# ZUGC-RNA degradation generates immunosuppressor to evade immune responses in eukaryotes

**DOI:** 10.1101/2025.01.27.633273

**Authors:** Shuliang Gao, Mengting Chen, Douglas Wich, Hanan Bloomer, Zhiyuan Qu, Huiwen Guan, Qiaobing Xu

## Abstract

Among the hundreds of modified nucleosides identified in terrestrial life, 2-amino-6-aminopurine (Z) is widely recognized as a prominent modified purine. Recently, RNA written with the ZUGC alphabet shows significant potential in RNA therapeutics as a synthetic biosystem. Here, we demonstrate that ZUGC-RNA can evade immune recognition in eukaryotes, independent of factors such as RNA length, sequence, 5’-triphosphate, modified uridine, and secondary structure. Notably, we discovered that both the degradation of ZUGC-RNA and metabolites of Z-nucleotides can function as immunosuppressors, silencing TLR7 sensing to block immune responses. This mechanism differs from that of pseudo-uridine (Ψ) modified RNA currently in use. ZUGC-RNAs also demonstrate broad applicability across multiple neural cell types. Our findings provide valuable insights for developing more tolerable RNA-based drugs and designing immunomodulators targeting TLR7. In addition to the potential prebiotic relevance of Z, our finding not only contributes to understanding the RNA world hypothesis but also provides new insights into the exploration of the origin of life.

## Introduction

Natural evolution has shaped the diversity of modified bases in ribonucleic acid (RNA) (*1*). Among these, 2-amino-6-aminopurine (Z) is a purine analog thought to have coexisted in the RNA world during early Earth with 6-aminopurine (A), 2-amino-6-oxopurine (G), cytosine (C), and uracil (U) (*1–4*). The Z base is not only found in terrestrial life forms but also in meteorites (*5, 6*). In the RNA world hypothesis, Z is considered a modified base that may have played a significant role in the origin of early life. In recent years, researchers have discovered that the biosynthetic pathway for Z is widespread (*7*). For example, cyanobacteria, which have existed on Earth for 3.5 billion years, host cyanophages like S-2L that possess a deoxyribonucleic acid (DNA) genome where A is completely replaced by Z. Although there is no direct evidence of Z base involvement in prokaryotic immune processes, the emergence of Z-genomes is thought to help phages evade host restriction endonuclease attacks on their DNA (*8, 9*). The discovery of Z biosynthetic pathways has provided new insights into synthetic life forms. As a fundamental principle of biochemistry, nucleotides can be synthesized *de novo* or recycled through salvage pathways in living organisms (*10*). This suggests the potential existence of ribonucleotides and Z-incorporated RNA. While naturally occurring Z-incorporated RNA has not yet been identified, considerable evidence supports its possibility. For instance, Z-incorporated RNA and DNA have been observed when prokaryotes and eukaryotes are supplemented with Z (*11*). Moreover, RNA polymerases from T7 bacteriophage and *Salmonella typhimurium* SP6 phage exhibit significant activity in synthesizing ZUGC RNA (Z-RNA) during *in vitro* transcription (IVT) (*12, 13*). With the rise of nucleic acid therapeutics, research on the biological applications of Z is increasing, including its use in messenger RNA (mRNA) and small RNAs, such as guide RNA (gRNA) for genome editing (*2, 12, 14, 15*).

To date, over 150 types of modifications have been identified in cellular RNA (*16–18*). However, the modified nucleobases currently used in mRNA therapeutics are primarily pyrimidines (e.g., pseudo-uridine, Ψ, and 5-methylcytosine, m5C), which can enhance protein expression while reducing the immunogenicity of exogenous RNA. The cellular mechanisms underlying RNA sensing and immune response initiation are influenced by various factors of RNA, such as base modifications, purity, sequence, secondary structure, and length (*19–22*). These complexities leave the working mechanisms of Z-base or ZUGC nucleic acid systems on cellular immune responses largely unknown, even 80 years after their discovery (*23*). Eukaryotes contain numerous pattern recognition receptors (PRRs) that detect nucleic acids, including the Toll-like receptor (TLR) family (e.g., TLR7 and TLR8) and the retinoic acid-inducible gene I (RIG-I)-like receptor (RLR) family of RNA sensors (*20, 21, 24*). Understanding how Z-based nucleic acids interact with these immune pathways is critical for leveraging their therapeutic potential and expanding roles of modified purines in immune regulation.

In this study, we investigated the effects of Z-RNA nucleic acids on immune activation in mammalian cells. We found that Z-RNA nucleic acids can evade recognition by the mammalian immune system, including the TLR7 receptor and RIG-I receptor, thereby preventing inflammatory responses. This immune evasion is independent of factors such as RNA length, sequence, modified uridine, and secondary structure. Notably, we identified a previously unreported regulatory mechanism: Z-RNA degrades into derivatives that suppress TLR7 receptor responses to even AUGC-RNA (A-RNA), which is distinct from that of modified-uridine-dependent RNA (**Fig. 1**). These results suggest the existence of a unique mechanism of immune evasion from purine-modified RNA base. It is the first time to establish a connection between Z-based nucleic acids and specific immune sensors. Our study also provides valuable insights for designing TLR7-targeting immunomodulators and advancing RNA-based drug development and production.

**Fig. 1.**
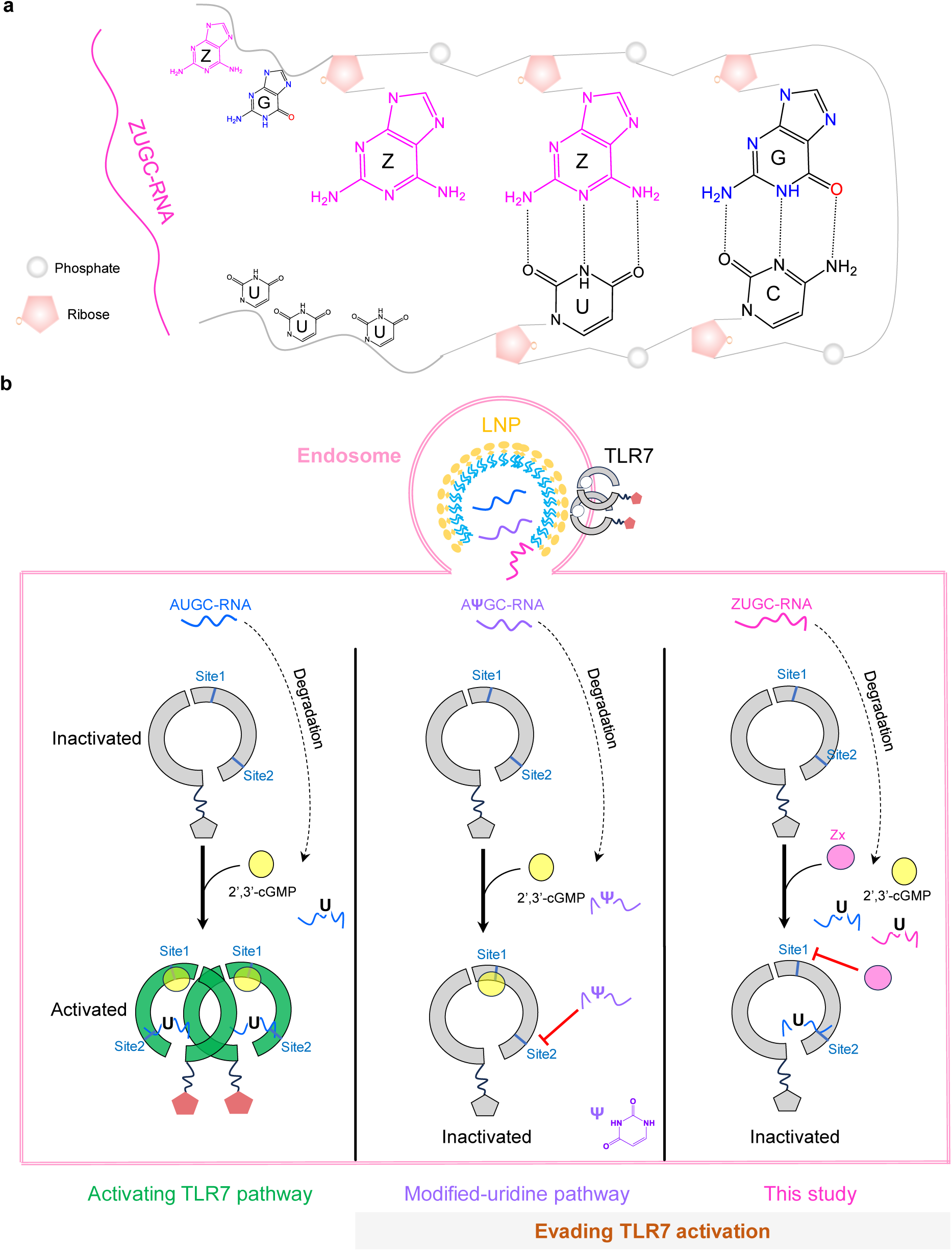
Highlights of this study. **(a)** Schematic illustration of single stranded ZUGC-RNA (Z- RNA). **(b)** Exogenous AUGC-RNA (A-RNA), upon entering the cell via the endosome, is degraded into products including 2’,3’-cGMP and oligonucleotides containing NUN motifs, which activate the TLR7 receptor by binding site 1 and site 2, respectively. This activation triggers downstream signaling pathways, leading to inflammatory responses including type-I interferon (IFN) production. Modified uridines (e.g., pseudo-uridine, Ψ), have been widely utilized to reduce the immunogenicity of exogenous RNA by suppressing RNA recognition. In this study, we identified an unreported immunomodulatory mechanism mediated by the Z base in the form of Z-RNA that enables TLR7 immune evasion independent of modified uridine.

## Results and discussion

### The integrity of Z-mRNA IVT transcripts is influenced by the 5’-UTR sequence

In our previous study (*12*), we observed that significant amounts of non-full-length transcripts were produced during *in vitro* transcription (IVT) reactions when synthesizing enhanced green fluorescent protein (EGFP) Z-RNA approximately 1000 nucleotides (nt) in length. We hypothesized that the stability of the secondary structure of transcripts might be contributing to this issue. To test whether optimizing the sequence to reduce stability could improve *in* vitro transcribed Z-RNA purity, we modified the 5’ untranslated region (5’-UTR), a critical element that influences mRNA secondary structure stability (*25*). We compared two 5’-UTR sequences: UTR1, a 66-nt sequence from a commercial plasmid (pcDNA3.1/Zeo(-)) previously used for Z-mRNA synthesis (*13*), and UTR2, a 206-nt sequence validated in our other work (*26*). UTR2 exhibited a more stable secondary structure than UTR1, as indicated by thermodynamic free energy values (- 56.60 kcal/mol vs. -12.68 kcal/mol) (**Fig. s1**), and led to greater stability of EGFP and Firefly luciferase (FLuc) transcripts (**Fig. s2 and s3**). However, UTR1 significantly improved the transcription quality of Z-RNA by reducing the generation of non-full-length transcripts (**Fig. s4**). Therefore, unless otherwise specified, subsequent experiments utilized the UTR1 sequence for mRNA synthesis (**Fig. 2a**).

**Fig. 2.**
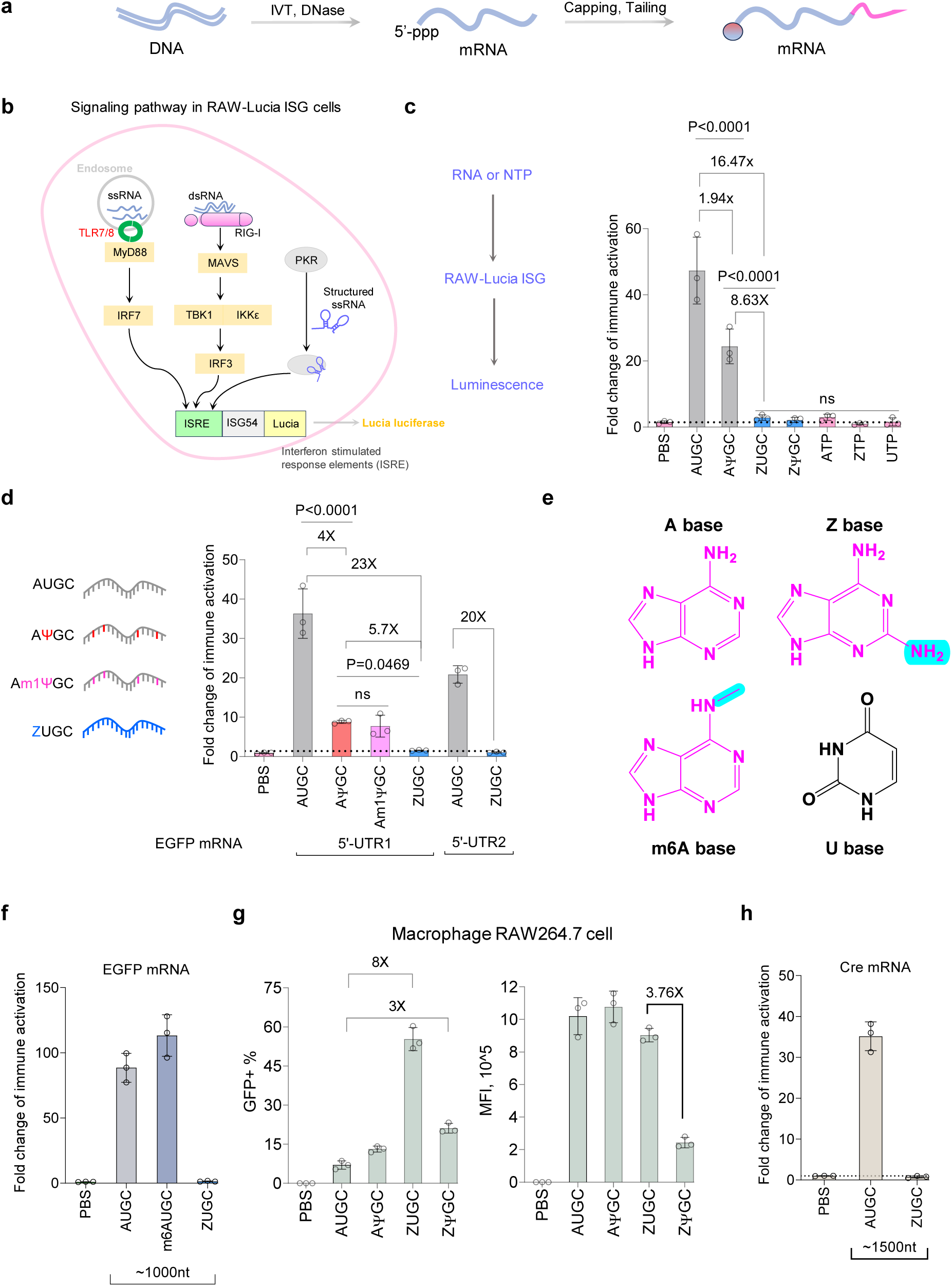
Z-mRNA evades immune activation in eukaryotes. (**a**) Workflow of preparing linear mRNA using *in vitro* transcription (IVT). Cap0 structure and poly(A) were used for capping and tailing, respectively. (**b**) Representative RNA sensing pathways for ssRNA and dsRNA in RAW-Lucia ISG cells. (**c**) Interferon (IFN) activation mediated by various mRNA constructs and NTPs (n = 3). (**d**) IFN activation mediated by mRNA constructs with N1-methyl-pseudouridine (Ψ) incorporation and the UTR2 sequence (n = 3). (**e**) Structures of A, Z, U and m6A bases. (**f**) IFN activation mediated by m6A-mRNA (n = 3). (**g**) EGFP expression in RAW264.7 cells (n = 3). (**h**) IFN activation induced by Cre Z-mRNA. The UTR1 sequence was used for this construct (n = 3). All data is presented as mean ± SD. For (c-d), statistical analysis was performed using an ordinary one-way ANOVA with Tukey’s multiple comparisons test. *P* values of <0.05 were considered statistically significant. ns, not significant.

**Fig. 3.**
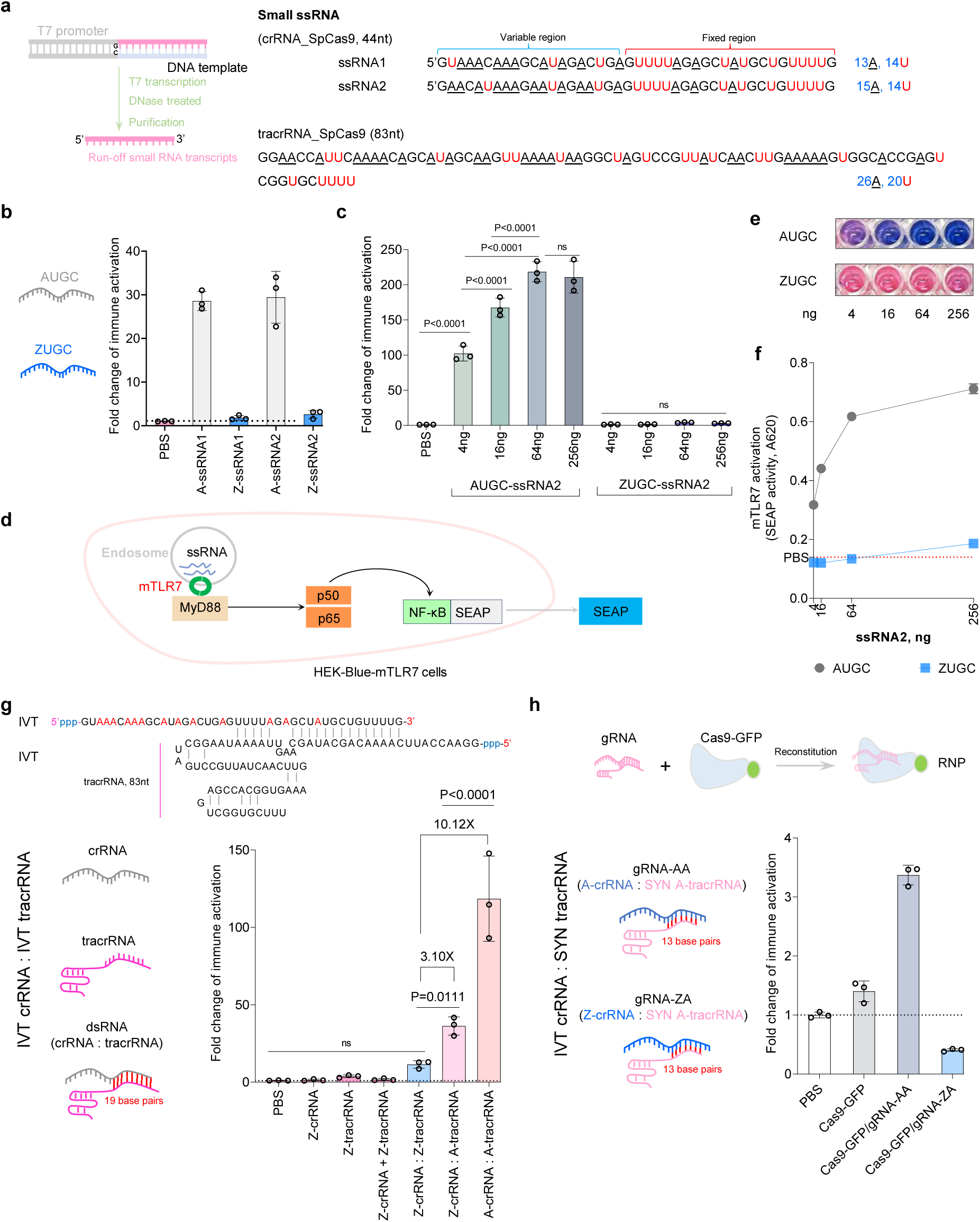
Both single and double stranded small RNAs written by ZUGC prevent immune response. (**a**) IVT and RNA sequences used in this analysis. A was completely replaced by Z in Z-RNA. (**b**) Immune activation mediated by Z-RNA in RAW-Lucia ISG reporter cells. A-ssRNA refers to AUGC-ssRNA and Z-ssRNA refers to ZUGC-ssRNA (n=3). (**c**) Immune activation at different doses of A-ssRNA and Z-ssRNA in ISG cells (n = 3). (**d**) Sensing pathway of mTLR7 in HEK-Blue-mTLR7 reporter cells. SEAP, secreted embryonic alkaline phosphatase. (**e**) Representative images of HEK-Blue-mTLR7 assay. (**f**) Quantification of HEK-Blue-mTLR7 assay (n = 3). (**g**) Immune activation induced by gRNA containing a dsRNA region in ISG cells (n = 3). “+” indicates combination of dual RNAs. “:” indicates annealing dual RNAs. (**h**) Immune activation is induced by ribonucleoproteins (RNPs). SYN tracrRNA refers to chemical synthesized tracrRNA. All data is presented as mean ± SD. For (c, h), statistical analysis was performed using an ordinary one-way ANOVA with Tukey’s multiple comparisons test. *P* values of <0.05 were considered statistically significant. ns, not significant.

### The immune evasion capability of Z-mRNA is independent of sequence, 5’-triphosphate, and modified uridine

RNA signaling pathways include receptors such as TLR7, which senses single-stranded RNA (ssRNA) in endosomes, and cytosolic sensors such as RIG-I and RNA-activated protein kinase (PKR). Once activated, these sensors trigger downstream interferon (IFN) responses (*24*). We employed the IFN-stimulated response element (ISRE)-reporter cell line RAW-Lucia ISG cells (ISG), derived from the murine macrophage RAW264.7 cell line, to assess immune responses. RAW264.7 cells can express multiple PRRs including TLR7 and RIG-I (*27*). In ISG cells, reporter luciferase activity correlates with the intensity of immune activation (**Fig. 2b**) (*28*). EGFP mRNA with different base compositions (AUGC, AΨGC, ZUGC, ZΨGC) was delivered to ISG cells using lipofection. As expected, AUGC-mRNA elicited a strong IFN response, while AΨGC-mRNA significantly reduced immune activation. Surprisingly, Z-RNA eliminated IFN immune activation, regardless of whether Ψ was incorporated. Like unmodified nucleoside 5’-triphosphates (NTPs) ATP and UTP, ZTP nucleotides did not trigger an IFN response (**Fig. 2c**). Furthermore, the immune evasion properties of Z-RNA were unaffected by the Cap0 structure or the poly(A) tail (**Fig. s5)**. Compared to Ψ, N1-Methyl-pseudouridine (m1Ψ) has been shown to more effectively reduce IVT mRNA innate immunogenicity (*22, 29*). We found that A(m1Ψ)GC-mRNA and AΨGC-mRNA mediated comparable levels of IFN responses, reducing immune activation by 4-fold compared to AUGC but still significantly higher than Z-RNA (**Fig. 2d**). Additionally, while UTR sequences influenced the immunogenicity of AUGC-mRNA, Z-RNA did not induce immune activation, regardless of whether UTR1 or UTR2 was used (**Fig. 2d**). These findings indicate that even non-full-length transcripts of EGFP Z-mRNA produced during IVT remain non-immunogenic.

The application of modified purines has faced challenges, such as N6-methyl-adenine (m6A) incorporation, which reduces immunogenicity but severely suppresses protein expression (*30, 31*). Z and m6A represent 2-amination and N6-methylation modifications of adenine, respectively (**Fig. 2e**). To compare the effects of Z and m6A on RNA immunogenicity, we tested m6AUGC-mRNA and Z-mRNA. Unlike Z-mRNA, m6AUGC-mRNA induced strong IFN responses comparable to AUGC-mRNA (**Fig. 2f**). This contrasts with previous reports that m6A incorporation could eliminate the immunogenicity of exogenous mRNA in various cell types (*19*). This discrepancy may be due to differences in m6AUGC-mRNA sensing pathways in murine macrophage cell lines used in this study or sequence-dependent effects of m6A on immunogenicity. Based on these findings, we did not pursue further comparative studies with m6A.

Next, we treated RAW264.7 macrophages with EGFP mRNA using lipofection. The low immunogenicity of Z-RNA likely contributed to the highest percentage of GFP-positive cells in macrophages (**Fig. 2g and Fig. s6**), approximately 8-fold higher than AUGC-mRNA. Notably, Z-RNA achieved the same GFP expression levels as AUGC and AΨGC. Immune responses induced by different RNAs in RAW264.7 cells were consistent with the ISG assay results, where Z-RNA exhibited decreased immune activation (**Fig. s7**). To determine whether the immunogenic properties of Z-RNA were dependent on sequence or length, we synthesized mRNA encoding Cre recombinase (1053 nt coding sequence, 333 nt longer than EGFP). Cre Z-mRNA also did not elicit immune responses (**Fig. 2h**) but exhibited detectable activities in Cre-loxP-reporter cell lines, including mouse Neuro2a cells (N2A) and human BJ fibroblast cells (BJ) (*32*). Interestingly, Cre Z-mRNA yielded a higher percentage of GFP-positive BJ fibroblasts (**Fig. s8**).

To assess whether the differences in IVT ssRNA immune activation were due to double-stranded (dsRNA) impurities, we performed immune dot blot analysis with J2 antibodies, which detect dsRNA longer than 40 nt without bias for modified RNA (*33*). The analysis revealed no detectable dsRNA impurities in AUGC, m6AUGC or ZUGC, whereas AΨGC and A(m1Ψ)GC led to low levels of dsRNA (**Fig. s9**). This suggests that the immunogenicity of transcripts with Ψ or m1Ψ may be attributed to dsRNA impurities. Additionally, these findings indicate that Z incorporation preserves IVT RNA purity by minimizing dsRNA impurities.

In mammalian cells, the 5’-triphosphate (5’-ppp) group of ssRNA is a key structural feature for activating RIG-I in the cytoplasm (*34, 35*). Additionally, structured ssRNA with a 5’-ppp group can activate PKR to induce immune responses (*24, 35*). The intrinsic 5’-ppp of IVT RNA presents safety concerns and limits its application in certain contexts (*36–38*), often necessitating the use of chemically synthesized RNA as an alternative (*34, 39, 40*). To evaluate the impact of 5’-ppp on Z- mRNA immunogenicity, we treated non-capped IVT mRNA with Antarctic phosphatase (AnP), an alkaline phosphatase, to remove the 5’-ppp group (*41*). As shown in **Fig. s10**, AnP treatment had no effect on the immunogenic profile of Z-mRNA, as both treated and untreated Z-mRNA failed to induce IFN responses. As expected, dephosphorylation significantly reduced the immunogenicity of AUGC- and AΨGC-RNA, consistent with previous studies.

In summary, immunomodulatory properties of ZUGC-mRNA are independent of pyrimidine modifications, length, 5’-ppp, and sequence in IVT RNA. Importantly, the contrast in immune activation capabilities between Z and m6A further highlights the need to investigate their underlying mechanisms.

### Z substitution enables small RNA to block TLR7 and RIG-I immune responses

Small RNA (e.g., siRNA, gRNA) occupies a critical position in RNA drug development (*42, 43*). To reduce immunogenicity, small RNA is often chemically synthesized. However, this approach frequently encounters challenges due to sequence dependency (*44, 45*). TLR7 and TLR8, two key receptors for sensing endosomal ssRNA, trigger immune responses based on NUN motifs or uridine (U) ligands (*46, 47*). While long mRNA can be optimized for U-depletion using synonymous codon substitution strategies (*48*), small RNA often retains GU-rich sequences, which enhance TLR7/8 recognition. To investigate the effect of Z incorporation on the immunogenicity of small RNA, we selected crRNA (44 nt) and tracrRNA (83 nt) sequences of *Streptococcus pyogenes* Cas9 (SpCas9) as examples. Aside from differences in sequence, tracrRNA is more prone to forming stem-loop secondary structures under physiological conditions compared to crRNA.

Two crRNA sequences, ssRNA1 and ssRNA2, which were previously validated for effective SpCas9 function, were used in this study (*12*). Both sequences feature 5’-end A-rich variable regions and 3’-end U-rich fixed regions (**Fig. 3a**). When IVT RNA was delivered via lipofection, both AUGC-ssRNA1 (A-ssRNA1) and AUGC-ssRNA2 (A-ssRNA1) induced strong IFN responses. In contrast, Z-ssRNA blocked immune activation for both sequences, reducing responses to near-baseline levels (**Fig. 3b**). Furthermore, varying doses of Z-ssRNA consistently failed to initiate immune activation (**Fig. 3c**). Interestingly, when A-ssRNA and Z-ssRNA were co-delivered, the intensity of immune activation positively correlated with the total RNA dose (**Fig. s11**). This suggests that Z-ssRNA may interact with A-ssRNA to induce immune activation.

Murine TLR7 (mTLR7), rather than TLR8, is the primary endosomal receptor in RAW264.7 macrophages for sensing GU-rich ssRNA (*47, 49, 50*). To investigate mTLR7 activation, we utilized HEK-Blue-mTLR7 reporter cells (*51*), which express the murine mTLR7 gene alongside an NF-κB/AP-1-inducible SEAP reporter gene (**Fig. 3d**). Lipofectamine transfections of A-ssRNA into these cells revealed an immune activation pattern similar to that observed in ISG cells. Z- ssRNA failed to activate mTLR7 at all tested doses (**Fig. 3e, 3f**). Long-chain ssRNA (EGFP ∼1000nt or FLuc ∼2000nt) exhibited the same characteristics in HEK-Blue-mTLR7 cells (**Fig. s12**). These results demonstrate that ssRNA composed of ZUGC evades mTLR7-mediated recognition.

RIG-I, a critical receptor for sensing cellular dsRNA (≥19 bp) and ssRNA, can also be evaded by incorporating modified bases into RNA (*34, 39, 52*). We annealed crRNA and tracrRNA components to form a full gRNA with a dsRNA region containing 19 base pairs (crRNA:tracrRNA, 13 A:U base pairs, 68.4% A:U content) and evaluated the effect of Z incorporation on immune activation. As shown in **Fig. 3g**, Z-tracrRNA, despite being twice as long as Z-crRNA, did not trigger immune activation. However, gRNA-ZZ (Z-crRNA:Z-tracrRNA) induced a slightly higher IFN response than ssRNAs, although not statistically significant. gRNA-ZA (Z-crRNA:A- tracrRNA), which included 3 Z:U base pairs (15.7% A:U content), elicited a significantly stronger immune response, increasing activation by 3.1-fold compared to gRNA-ZZ. As expected, standard gRNA-AA (A-crRNA:A-tracrRNA) elicited the strongest immune activation, which was 10.1-fold higher than gRNA-ZA and 31.3-fold higher than gRNA-ZZ. Notably, A-crRNA alone and gRNA- AA triggered the same level of immune activation at equivalent doses (**Fig. s13**). We further tested dsRNA with 70 base pairs and found that ZUGC-dsRNA reduced human RIG-I activation by 28.3- fold compared to AUGC-dsRNA (**Fig. s14**). Similar to modified uridines (*34*), Z-incorporated small ssRNA can also block RIG-I sensing. These findings align with the results from ISG cell testing, suggesting that small Z-dsRNA and Z-ssRNA can reduce RIG-I recognition in both murine and human systems.

Chemically synthesized AUGC-tracrRNA (SYN-tracrRNA), which lacks a 5’-ppp group and includes specific modifications, is often used to reduce immunogenicity and enhance gene editing efficiency (*53*). We compared IVT Z-RNA with two chemically synthesized RNAs in ISG cells and observed no significant differences in immune activation (**Fig. s15**). We then compared the immunogenicity of IVT-tracrRNA (83nt) to SYN-tracrRNA (67nt). Hybridized gRNA-AA containing SYN-tracrRNA significantly reduced immunogenicity by 6-fold (**Fig. s16a and b**). Notably, gRNA-ZA with SYN-tracrRNA failed to elicit an immune response, with levels comparable to phosphate buffered saline (PBS). These findings suggest that incorporating Z bases can eliminate gRNA immunogenicity without relying on complex chemical modifications.

To investigate the impact of Z-substitution on the immune activation of Cas9-gRNA ribonucleoproteins (RNPs), we reconstituted Cas9-GFP fusion proteins with different gRNAs (prepared by hybridizing IVT-crRNA with SYN A-tracrRNA). As shown in **Fig. s16c and s16d**, the delivery of RNPs *via* lipofection demonstrated that gRNA-ZA achieved similar transfection efficiency to gRNA-AA. Encouragingly, gRNA-ZA elicited no immune activation, whereas RNPs containing gRNA-AA triggered the highest IFN response (**Fig. 3h**). These findings indicate that the incorporation of Z bases can minimize the immunogenicity of Cas9 RNPs.

In summary, Z-incorporation enables small RNA to effectively evade PRR-mediated immune recognition in macrophages, regardless of unmodified uridine content. Specifically, Z-RNA bypasses recognition by mTLR7 and RIG-I receptors, thereby preventing downstream IFN responses.

### Z-RNA evades immune response regardless of hairpin structure and unmodified uridine content

Encouraged by the results above, we sought to explore whether ZUGC exhibits a unique mechanism for evading immune responses, as Z-RNA lacks modified uridine. We selected Cas12a crRNA sequences as the ssRNA for this study. Both our group and others have validated that Z- crRNA can maintain or enhance Cas12a cleavage and gene editing activity (*12, 15*), suggesting that Z-crRNA retains an accurate secondary structure. Unlike that of SpCas9, Cas12a crRNA sequences tend to maintain simpler secondary structures with a single stem loop under physiological conditions (**Fig. s17 and s18**). Cas12a crRNA sequences contain a fixed 5’-end region and a variable 3’-end region (**Fig. 4a**). We first prepared three crRNAs (crRNA1, 2, 3) via IVT and delivered them to ISG cells using lipofection. All A-crRNAs triggered strong IFN responses, while all Z-crRNAs prevented immune activation (**Fig. 4b**). To assess whether RNA delivered by lipofection induces TNF-α production in macrophages, we treated RAW264.7 cells with crRNAs and used their supernatants to stimulate HEK-Blue-mTLR7 reporter cells equipped with a TNF-α receptor. A-crRNAs led to increased TNF-α receptor activation compared to Z- crRNAs (**Fig. s19**). These results demonstrate that Z incorporation enables short ssRNA to evade immune responses regardless of small hairpin secondary structures. Additionally, Z-RNA prevents the activation of multiple cytokine pathways, including IFN and TNF. Removal of the 5’-ppp group did not alter the immunogenicity of Z-crRNA, consistent with observations from long ssRNA studies (**Fig. 4c**). While m6A modifications reduced crRNA immunogenicity, they still triggered a strong IFN response (**Fig. 4d**), further illustrating that those modifications at different purine positions have distinct impacts on the immunogenicity of small ssRNA.

**Fig. 4.**
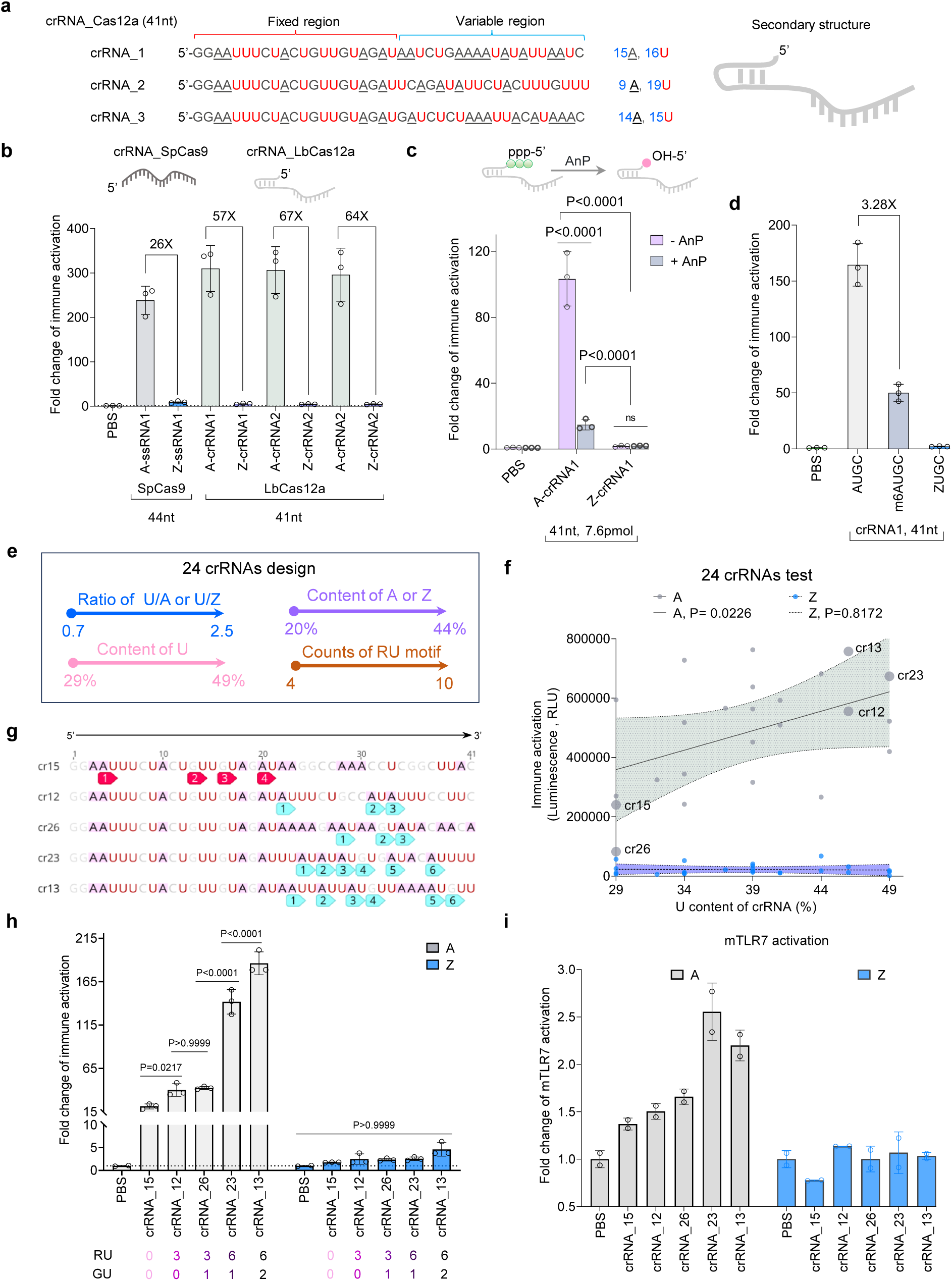
Z-ssRNA evades immune activation regardless of unmodified uridine content. (**a**) Cas12a crRNA sequences and secondary structure. (**b**) Immune activation of Cas12a crRNA in ISG cells (n = 3). (**c**) Effects of 5’-ppp group in IVT crRNA on immune activation in ISG cells (n = 3). (**d**) Immunogenicity of m6A and Z substituted crRNA in ISG cells. (**e**) Design of 24 crRNAs. (**f**) Immunogenicity of 24 crRNAs with various U content in ISG cells. Linear regression was analyzed using GraphPad. Samples cr12, cr13, cr15, cr23, and cr26 were highlighted by dots in larger size. (**g**) Representative sequences of the 5 selected crRNAs. Red numbered frames indicate RU motifs in the fixed region, and teal frames indicate RU motifs in the variable region. R = A or G. (**h**) Immune activation induced by the 5 crRNAs in (g) using ISG cells (n = 3). Counts of RU or GU motifs in each RNA sequence are presented at the bottom. (**i**) mTLR7 activation induced by the 5 crRNAs in (g) using HEK-Blue-mTLR7 cells (n = 2). All data is presented as mean ± SD. For (c), statistical analysis was performed using a 2way ANOVA with Sidak’s multiple comparisons test. For (h), statistical analysis was performed using an ordinary one-way ANOVA with Dunnett’s multiple comparisons test. *P* values of <0.05 were considered statistically significant. ns, not significant.

crRNA2 has a U content of 46.3%, making it 2.1 times the A or Z content (U/A ratio = 2.1). This high U content did not affect the immunogenicity of Z-crRNA, leading us to hypothesize that the immune evasion mechanism of Z-RNA differs from previously reported modified-uridine-dependent mechanisms. To test this hypothesis, we designed 24 crRNA sequences (cr4 to cr27) of identical length by modifying the 3’-end variable region sequences. These sequences had U/A (or U/Z) ratios ranging from 0.7 to 2.5, U content ranging from 29% to 49%, A or Z content ranging from 20% to 44%, and the number of RU motifs ranging from 4 to 10, where R represents purines (**Fig. 4e and s20**). Immune activation tests with these crRNAs revealed that all A-crRNAs induced strong IFN responses in ISG cells, with activation intensity positively correlating with U content and RU motifs counts but not with the U/A ratio (**Fig 4f, Fig. s21**). Encouragingly, all Z-crRNAs showed minimal and uniform immunogenicity. We selected five representative crRNAs (cr12, cr13, cr15, cr23, cr26) (**Fig. 4g**) for further study. The IFN activation strength induced by these A-crRNAs in ISG cells strongly correlated with the number of RU motifs (**Fig. 4h**), consistent with the initial screening results (**Fig. 4f**). We reasoned that ssRNA with high RU motif counts may be more prone to degradation, as lysosomal RNase T2 specifically cleaves RU motifs in ssRNA (*46*). In contrast, Z-crRNAs showed no significant difference from PBS controls. Additionally, these five crRNAs exhibited similar trends for mTLR7 activation as observed in ISG cells (**Fig. 4i**). These findings demonstrate that Z-RNA evades immune recognition, even with high U content.

To further test whether varying levels of Z substitution in IVT RNA (AZUGC-RNA) can influence immunogenicity, we prepared mRNAs (EGFP and FLuc) with varying ratios of Z substitution (A:Z ratios of 80:20, 50:50, and 20:80). As shown in **Fig. s22**, Z substitution at levels equal to or greater than 50% resulted in decreased immune activation and completely blocked mTLR7 activation, regardless of sequence composition. These results also indicate that the contents of A and Z can alter the immunogenic characteristics of AZUGC-RNA. This finding is similar to that of other modified pyrimidines (*19*).

In summary, these results indicate that Z-RNA immunogenicity is unaffected by U content, further supporting our hypothesis of the unique immune evasion mechanism of Z based RNA.

### Both Z-RNA degradation and metabolites of Z nucleotides enable blocking mTLR7 stimulation by A-RNA

As previously reported (*46, 47, 54, 55*), lysosomal RNases, particularly RNase T2, mediate the degradation of endosomal ssRNA in coordination with phospholipase exonucleases (PLDE) to produce ligands for TLR7 activation. These ligands, which include 2’,3’-cGMP or guanosine molecules and pyrimidine-rich oligonucleotides (typically containing NUN motifs, where N can be A, U, G, or C, with a pyrimidine preference), bind to TLR7 at site 1 and site 2, respectively (**Fig. 5a**). Binding these ligands induces TLR7 dimerization, trigger downstream signaling pathways and activating IFN responses. While modified nucleotides can enhance RNA resistance to RNase cleavage (e.g., Ψ improves RNA resistance to RNase L digestion), their effect on delaying degradation is minimal (*55, 56*), as endosomal environments exhibit high levels of diverse RNases (*57, 58*). It was previously demonstrated that Z-RNA does not exhibit improved resistance to RNase cleavage (*12, 13*) . This suggests that RNase resistance is not the primary reason for the immune evasion properties of Z-RNA. Additionally, the degradation of EGFP Z-mRNA still generates NUN ligands capable of activating TLR7 site 2 (**Fig. S23**). These analyses motivate us to focus on TLR7 site 1. We analyzed the structural interactions between TLR7 site 1 and its ligand, 2’,3’-cGMP. Structurally, guanine (G) and 2,6-diaminopurine (Z) are highly similar, differing only at the 6-position (oxo versus amino groups). Structural modeling suggests that TLR7 site 1 can bind Z-derived molecules (**Fig. 5b and 5c**). Compared to 2’,3’-cGMP, the analog 2’,3’-cAMP is a weaker ligand for TLR7 site 1 (*54*), which suggests that both 2-NH2 and 6-NH2 generate crucial binding ability with TLR7 residues. Furthermore, many small-molecule TLR7 antagonists share a purine ring as their core structure and indicate the critical roles of 6-amino group binding to TLR7 (*59–62*). Similar to G, Z have the basis to form strong hydrogen bonds with nearby residues including K432 and D555 in TLR site 1. However, the unique electronic distribution of Z’s diamino group significantly alters the strength and distribution of these hydrogen bonds, potentially interfering with TLR7 recognition and activation.

**Fig. 5.**
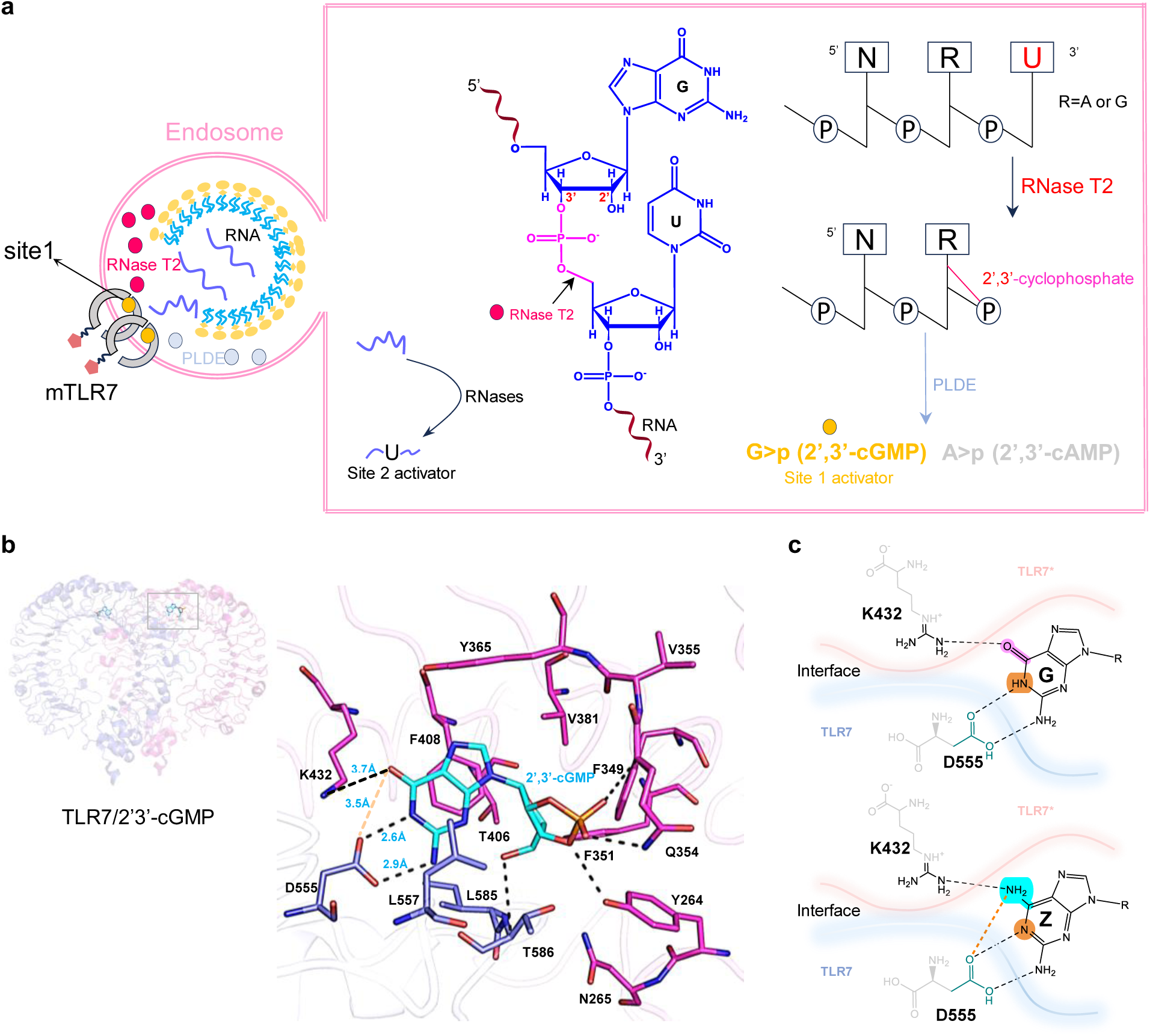
Z-RNA degradation may generate cZMP to suppress TLR7 activation. (**a**) Previously reported mechanism of endosomal RNA degradation. PLDE, phospholipase exonucleases. (**b**) Detailed view of 2’3’-cGMP binding at site 1 of TLR7 (PDB: 6IF5). The carbon atoms of 2’3’-cGMP are colored sky blue. Dashed line represents hydrogen bonds or distances between two groups. (**c**) Interaction between G or Z and D555. Dashed lines represent hydrogel bonds, with the weight of the dashed lines representing the strength of the hydrogen bond interaction. Single-letter abbreviations for the amino acid residues are as follows: A, Alanine; C, Cysteine; D, Aspartic acid; F, Phenylalanine; K, Lysine; L, Leucine; S, Serine; T, Threonine; V, Valine; Q, Glutamine; N, Asparagine; Y, Tyrosine.

One possible compound from Z-RNA degradation that may interact with TLR7 is the cyclic nucleotide (cNMP) form of Z (cZMP), which could affect TLR7 activation (**Fig. 6a**). Here, we propose naming this compound Zx. Cyclic purine nucleotides, such as cyclic adenosine monophosphate (cAMP) and cyclic guanosine monophosphate (cGMP), are well-established second messenger molecules in higher eukaryotes (*63, 64*). Both RNases and nucleotidyl cyclases (NCs) are capable of converting RNA and 5’-nucleoside triphosphates (NTPs) into cyclic nucleotides (cNMP) (*64–66*).

**Fig. 6.**
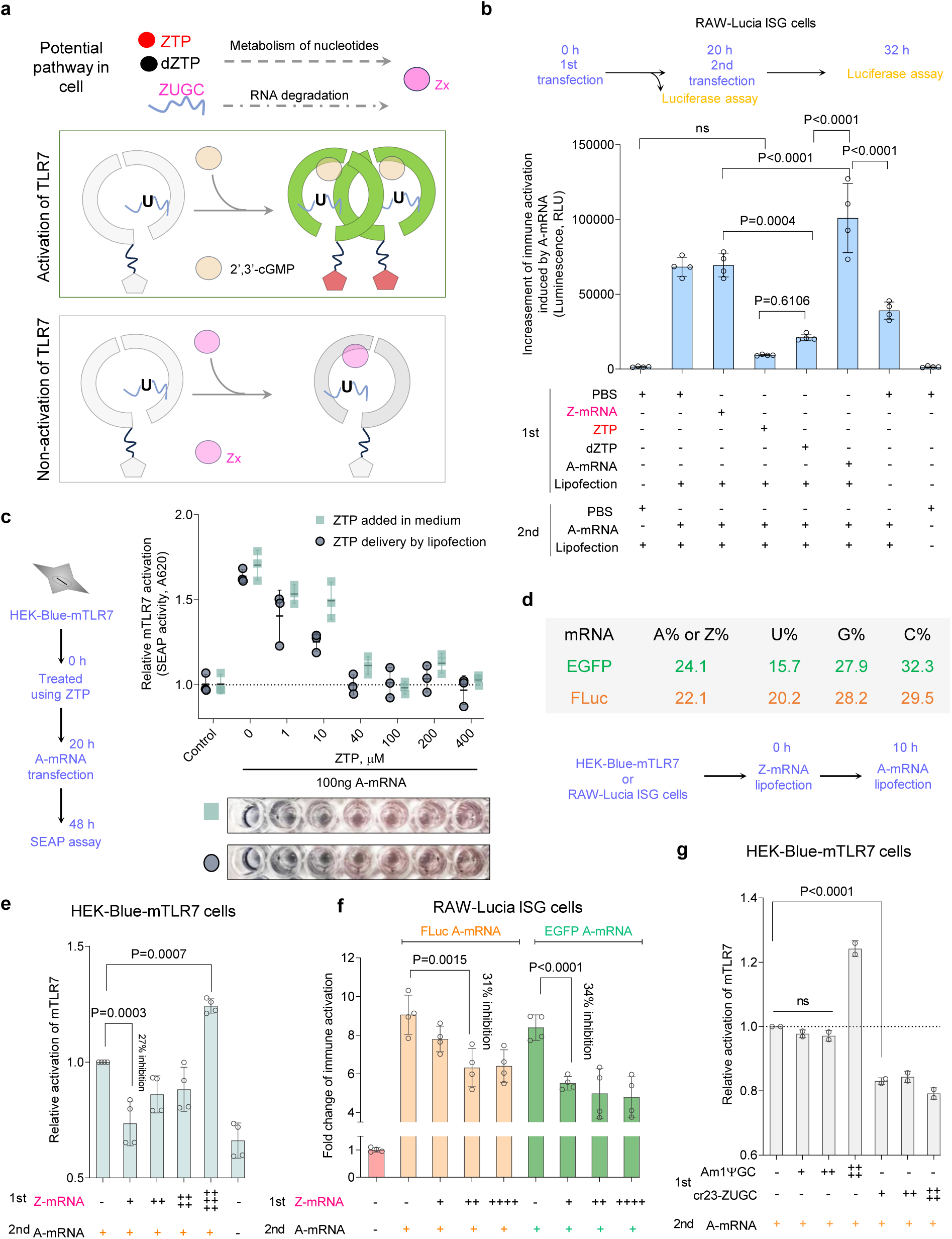
Z-RNA degradation functions as immunosuppressor to modulate TLR7 activation. (**a**) Schematic representation of the TLR7 activation and inhibition. (**b**) Immune activation induced by A-mRNA and with the addition of Z inclusion products in ISG cells (n = 4). For ZTP and dZTP, 400 µM dose was used. For A-mRNA and Z-mRNA, 100 ng Cre mRNA was used. (**c**) mTLR7 activation with increasing concentrations of ZTP (n = 3). The x-axis indicates experimental dose of ZTP. The conditions are divided into two stages (1st and 2nd). Representative well-images of assay are presented at the bottom. (**d**) Details of mRNAs used for (e) and (f). (**e**) mTLR7 activation induced by Z-mRNA or A-mRNA in HEK-Blue-mTLR7 cells (n = 4). (**f**) mTLR7 activation induced by Z-RNA or Am1ΨGC-RNA in HEK-Blue-mTLR7 cells. The Am1ΨGC-RNA is same to that of Fig. The cr23-ZUGC is same as Z-crRNA23 in Fig.4h. (**g**) Immune activation of A-RNA or Z-RNA in ISG cells (n = 4). All data is presented as mean ± SD. For (b, d, e, f, g), the conditions are divided into two stages (1st and 2nd). Positive (+) or negative (-) signs represent the presence or absence of specific treatments. For (e, f, g), the positive (+) signs labeled in black also indicate the dose (50ng) of the first treatment using EGFP mRNA, the (+) signs labeled in orange and green indicate the dose (30ng) FLuc and EGFP mRNA, respectively. For (b), statistical analysis was performed using a 2-way ANOVA with Sidak’s multiple comparisons test. For (e, f, g), statistical analysis was performed using an ordinary one-way ANOVA with Dunnett’s multiple comparisons test. *P* values of <0.05 were considered statistically significant. ns, not significant.

To test our hypothesis regarding Z-derived metabolites, we first examined whether nucleoside triphosphates, including pseudo-UTP, GTP, ZTP, 2-thio-UTP, and 5-methoxy-UTP, could inhibit immune responses to standard AUGC-RNA in ISG cells. Both ZTP and two modified UTPs (2- thio-UTP and 5-methoxy-UTP) significantly reduced the immune response to AUGC-RNA at a concentration of 400 µM, whereas pseudo-UTP and GTP did not exhibit this effect (**Fig. s24**). Given the characteristics of nucleotide salvage pathways (*67*), ZTP and dZTP are expected to produce similar metabolic products. We pretreated cells with ZTP and dZTP and observed that both significantly reduced the immune response induced by AUGC-RNA. Notably, ZTP demonstrated slightly stronger inhibitory effects than dZTP, bringing immune responses closer to baseline levels (**Fig. 6b and s25**). However, Z-mRNA pretreatment did not inhibit the immune response induced by AUGC-RNA, similar to the co-delivery of two RNAs (**Fig. s11**). This suggests that the inhibitory effect of Z metabolism on immune activation induced by AUGC-RNA may be dose-dependent. Next, we found that mTLR7 activation by AUGC-RNA was completely suppressed when cells were pretreated with 40 µM ZTP (**Fig. 6c**). This effect was consistent across both direct addition and lipofection treatments. Interestingly, this effective dose was 10 times lower than the concentration used in the ISG cells, likely due to differences in mTLR7 protein expression levels and PRR abundance between the two cell types (*19*). Neither GTP nor ATP alleviated the suppression caused by ZTP (**Fig. s26**). These results strongly suggest that certain metabolites of ZTP are responsible for inhibiting mTLR7 responses to AUGC-RNA. To further investigate this phenomenon, we examined whether Z-RNA pretreatment could inhibit immune responses to lower dose of AUGC-RNA (**Fig. 6d**). Lipofection of EGFP Z-mRNA at different doses into mTLR7 reporter cells resulted in a 27% inhibition of immune responses induced by A-mRNA (**Fig. 6e**). However, at high doses, Z-RNA synergized with A-RNA to enhance TLR7 activation. We speculate that Z derivatives and G derivatives may competitively bind to TLR7 site 1. In ISG cells, Z-mRNA also functioned as an immunosuppressor, regardless of RNA sequences (**Fig. 6e and 6f**). However, modified uridine mRNA didn’t show suppression on immunogenicity of AUGC-RNA (**Fig. 6g**). Guanosine has been identified as an activator of TLR7 (*54*), therefore, we further investigated whether the nucleoside 2-amino-adenosine (Zr) could function as an inhibitor to

reduce TLR7 activation (**Fig. 7a**). As expected, co-delivery of Zr suppressed TLR7 immune activation induced by AUGC-RNA (**Fig. 7b and 7c**).

**Fig. 7.**
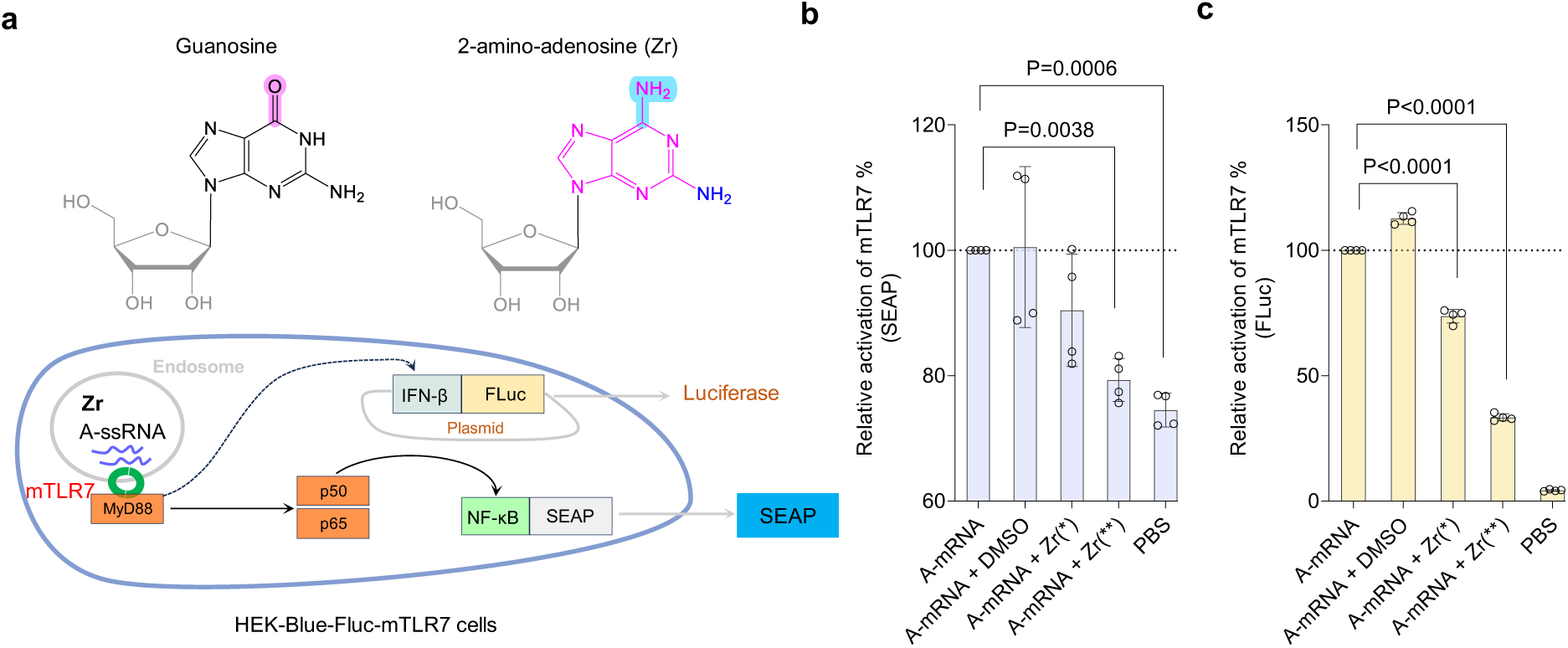
2-amino-adenosine (Zr) suppresses immune activation induced by AUGC-RNA. (**a**) Reporter pathway of mTLR7 activation used in this evaluation. (**b, c**) Zr can reduce the mTLR7 activation induced by A-mRNA. All data is presented as mean ± SD (n = 4). Statistical analysis was performed using an ordinary one-way ANOVA with Dunnett’s multiple comparisons test. P values of <0.05 were considered statistically significant. Positive (+) signs represent the superposition of two conditions of treatment. Asterisk (*) signs represent the dose of Zr.

In summary, we demonstrate that ZTP, Zr and Z-mRNA can exhibit immunosuppression on TLR7 sensing pathway. Although the exact identity of Z-derived metabolite Zx binding with TLR7 protein remains unclear, to the best of our knowledge, there have been no prior reports indicating that intracellular modified-purine-nucleotides and their metabolic products can function as immunosuppressors to inhibit TLR7 immune responses to exogenous RNA.

### Z-mRNA enables protein expression in multiple cell types

Next, we evaluated the compatibility of Z with other modified nucleotides for IVT and mRNA translation. Phosphorothioate modifications have been reported to enhance stability and translation in a cell free system (*68*). For comparison, we incorporated two additional modifications, GTPαS and CTPαS (**Fig. s27a**). However, both GTPαS and CTPαS resulted in considerable production of non-full-length transcripts during IVT (**Fig. s27b**) and drastically decreased mRNA yields (**Fig. s27c**). We evaluated the translation of various mRNAs in HEK293T and HeLa cells and found that phosphorothioate modifications completely blocked translation of EGFP mRNA in both cell lines (**Fig. s27d and e**). Notably, GTPαS and CTPαS also increase the immunogenicity of Z-mRNA (**Fig. s28**).

To test the general applicability of Z-mRNA for protein expression, we transfected EGFP Z-mRNA into 12 cell types including 9 non-cancer cell lines and 3 cancer cell lines. Among the 9 non-cancer cell lines, 6 were neural cell lines, such as human induced neural stem cells (iNSCs), human astrocytes (Astro), human brain vascular pericytes (Peri), human fetal brain-derived primary microglia (HMC3), brain microvascular endothelial cells (BMECs) and N2A cells (**Fig. s29-31**). In these neural cell types, Z-mRNA demonstrated similar or higher EGFP transfection efficiency and expression compared to A-mRNA (ranging from 2.9- to 12.9-fold). Similar results were observed in human umbilical vein endothelial cells (HUVECs), mouse dendritic cell line (DC2.4), human BJ fibroblast cells, and the Tsc2-null cell line TTJ. A-mRNA and Z-mRNA exhibited comparable efficacy in the mouse melanoma cell line B16F10, while Z-mRNA achieved higher EGFP expression than A-mRNA in the hepatocellular carcinoma cell line HepG2/C3A. These differences may be due to the variable expression of PRRs and immune-related activities across these cell types (*69, 70*). For instance, microglia cells are highly sensitive to TLR7 stimulation (*71*). These results demonstrate the generality and vast potential of Z-mRNA in various cells.

## Discussion

In this study, we identified that both ssRNA and dsRNA in the ZUGC form can evade immune recognition. This low immunogenicity is independent of sequence, secondary structure, length, and the presence of modified uridine. The enhanced binding strength of Z:U, which is 20-fold greater than A:U due to the introduction of an additional hydrogen bond (*2*), likely contributes to the immune evasion properties of Z-dsRNA. Geometric changes in the dsRNA structure caused by Z incorporation may prevent RIG-I recognition, a mechanism similar to that of other modified bases (*52*). In ISG cells, immune responses to AUGC-mRNA are primarily mediated by PKR signaling, suggesting that structural alterations in ZUGC-mRNA also enable its evasion of PKR detection (**Fig. s32**) (*56*). For ssRNA, Z-incorporation adopts a previously unreported immune evasion mechanism, where its degradation products act as immunosuppressors to inhibit TLR7 activation, even in the presence of immunogenic AUGC-ssRNA (**Fig.1**). While the precise structure of the immunosuppressor, tentatively named Zx, remains unidentified, our findings suggest it is likely cZMP according to the reported mechanism of endosomal RNA degradation (*46*). Eighty years ago, Z bases were observed to inhibit the replication of various viruses (*23, 72, 73*). This effect may partly arise from Z bases or their intracellular metabolites (e.g., Zx) suppressing TLR7 activation, thereby reducing excessive immune responses, mitigating cellular stress, and enhancing antiviral defenses (*74*). Our study is the first to establish a connection between Z-based nucleic acids and specific immune sensors, highlighting the potential broader immunomodulatory roles of Z bases during the propagation of ZTGC-based life forms.

Overexpression or abnormal activation of TLR7 signaling is well-documented to be associated with numerous autoimmune diseases and cancers, including systemic lupus erythematosus (SLE), type 1 diabetes (T1D), Sjögren’s syndrome (SS), rheumatoid arthritis (RA), lung cancer, and pancreatic cancer (*74, 75*). Safety concerns regarding RNA-based therapeutics in patients with autoimmune diseases persist (*76, 77*), as repeated high-dose administration of RNA drugs may lead to excessive TLR7 activation. Z-RNA may offer promising therapeutic strategies for these conditions. Further research into Zx and ZUGC-RNA will provide valuable insights and may inform the future design of safer and more effective RNA-based therapeutics.

We are currently unaware of any reports indicating that degradation products of RNA containing modified bases can inhibit TLR7 activation. Given the widespread presence of purine-modified bases in nature (*16, 18*), similar immune evasion mechanisms may be more prevalent than previously recognized. Although the pseudouridylation mechanism of uridine in RNA, involving the rearrangement of the imino group on the pyrimidine ring, is already well understood (*78, 79*), aminated modifications of purines have not been discovered in nature RNA to date. Due to the presence of the de novo synthesis pathway for the Z base in bacteria, it may differ from many pyrimidine modifications of RNA that occur post-transcriptionally. The immunoregulatory function of Z-RNA suggests the vast potential of this direction. Pathogens are known to deliver nucleic acids into host cells via vesicles to facilitate infection (*80*), and purine base modifications may enhance virulence by aiding in immune evasion (**Fig. 8**). Considering the anti-defense advantages of Z-modification, it is plausible that Z-modifications occur naturally, providing evolutionary benefits to certain pathogens. Numerous nucleotide derivatives are involved as messenger molecules in signaling pathways in bacterial defense, such as Cas10 protein from Type III CRISPR system sensing phage RNA can catalyze ATP and S-adenosyl methionine (SAM) to generate SMP-AMP for antiviral signaling (*81*). ZTP may also function similar roles. Additionally, there is long-standing speculation that Z-containing double-stranded DNA (Z-dsDNA) may also possess immunomodulatory effects. Our findings lay the groundwork for further investigations into the immune regulatory roles of Z, potentially uncovering new biological implications and therapeutic opportunities.

**Fig. 8.**
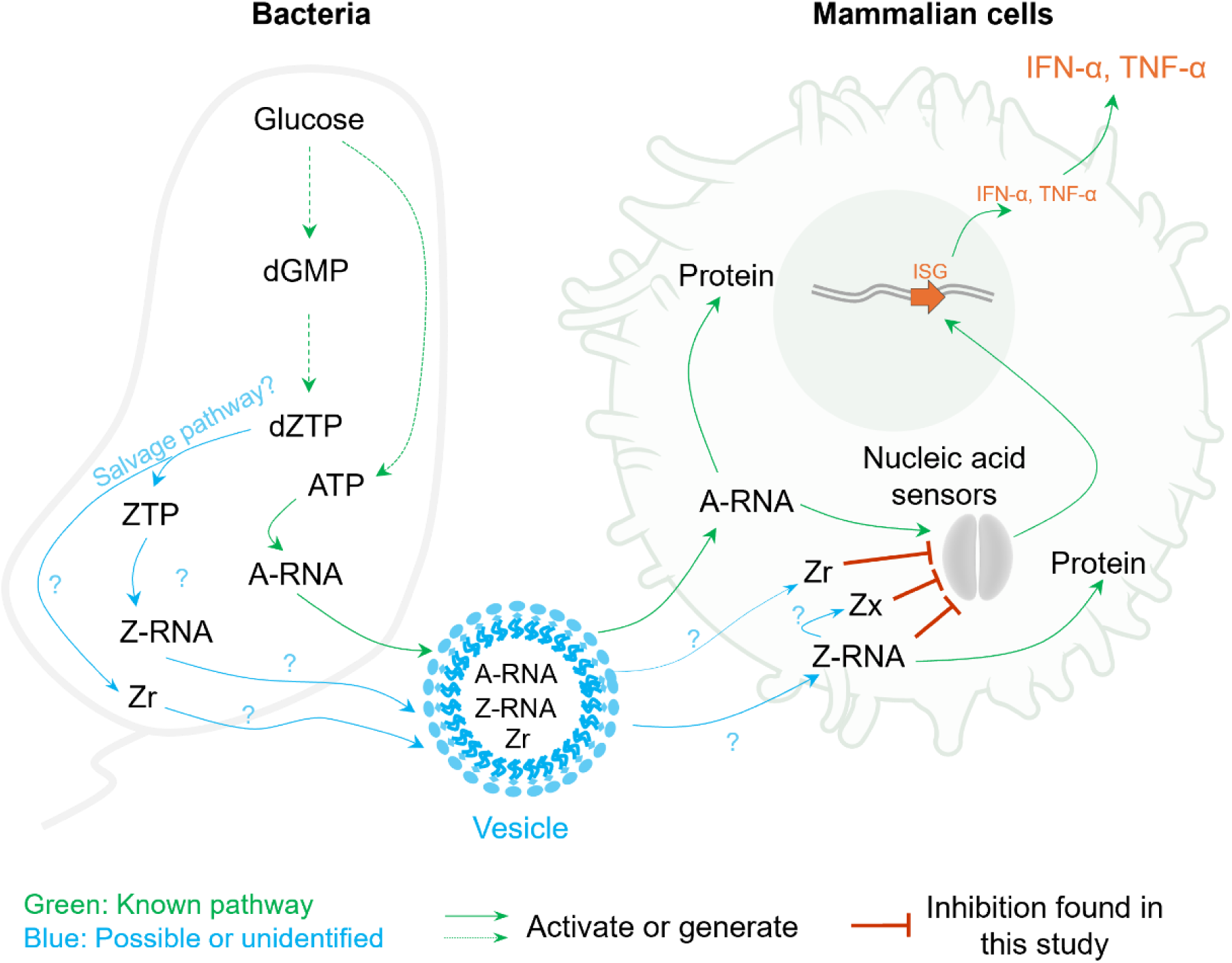
Proposed model of interaction between bacteria and mammalian cells mediated by Z base. Pathways in green and red color have been identified by previous research and our study. Pathways in blue are unknown to date.

## Acknowledgements

We thank Prof. Feng Zhang from Broad Institute of MIT and Havard for gifts of N2A-LoxP and BJ-LoxP cells. We thank Prof. Jinjun Shi from Harvard Medical School for the gift of HEK-Blue- mTLR7 cells. We thank Liam Power, MD/PhD student from Tufts University in Prof. David Kaplan’s lab, for plating neural cells.

## Author Contributions

Conceptualization: S.G. and Q.X. Funding acquisition: Q. X.

S.G. designed this work.

S.G., M. C., D. W. and H. G. performed the experiments.

1. Z. Q. performed TLR7 modeling.

S.G. and Q.X. analyzed the data.

S.G. wrote the manuscript. S. G., D.W., H. B., and Q. X. revised the manuscript.

## Data and Materials availability

All data associated with this study are present in the paper and/or the Supplementary Materials.

## Supplementary Materials

Materials and Methods Figs. S1 to S33

Tables S1 to S5 References (*81–82*)

