## Supplementary Material for "ZUGC-RNA degradation generates immunosuppressor to evade immune responses in eukaryotes"

Shuliang Gao, *et al*.

**The PDF file includes:**

Materials and Methods

Figs. s1 to s32

Tables s1 to s5

References *81-82*

#### Materials and methods

##### Construction of plasmids

Plasmid templates were constructed using standard cloning techniques, including digestion with a variety of restriction endonucleases and ligation using the Quick Ligation Kit (New England Biolabs, M2200S). DNA cloning and plasmid amplification were performed using One Shot Max efficiency DH5a-T1R Competent Cells (Thermo Fisher, 12297016). RNA or DNA oligonucleotides were synthesized by IDT and Genscript.

##### Cell culture

HEK293T cells were cultured as previous described (*80*). N2A, N2A-LoxP, BJ and BJ-LoxP cells were cultured as previous described (*81*). RAW264.7 cells (ATCC TIB-71) were cultured in DMEM supplemented with 10% fetal bovine serum (FBS). RAW-Lucia ISG cells (InvivoGen) and HEK-Blue-mTLR7 cells (InvivoGen) were cultured in complete DMEM containing 4.5 g/L glucose, 2 mM L-glutamine, 10% heat-inactivated FBS (incubated at 30 min at 56°C), 100 µg/mL Normocin, and Penicillin-Streptomycin (100 U/mL - 100 µg/mL). HeLa, B16F10, HepG2/C3A and Tsc2-deficient TTJ cells were cultured in DMEM (Sigma, D6429) supplemented with 10% FBS and 1% Gibco™ Penicillin-Streptomycin (10,000 U/mL). DC2.4 cells were cultured in RPMI-1640 (Sigma, R0883) supplemented with 10% FBS (ES-009-B), 1X L-glutamine (TMS-002-C), 1X non-essential amino acids (TMS-001-C), 1X HEPES Buffer Solution (TMS003-C) and 0.0054X β-Mercaptoethanol (ES-007E). Human induced neural stem cells (iNSCs) were cultured in NSC medium containing N3 medium and supplemented with 10 ng/mL recombinant mouse EGF (R&D systems) and 10 ng/mL recombinant human bFGF (R&D Systems), with or without 2 ng/mL doxycycline (Sigma). The N3 medium contains DMEM/F12 (Life Technologies) with 1 × Pen/Strep, 25 µg/mL insulin (Sigma), 50 µg/mL Apo-transferrin (Sigma), 1.28 ng/mL progesterone (Sigma), 16 ng/mL putrescine (Sigma), and 0.52 µg/mL sodium selenite (Sigma). Astrocytes and Pericytes were cultured in Astrocyte Medium (AM, Cat. #1801) and Pericyte Medium (PM, Cat. #1201), respectively. HMC3 cells (ATCC CRL-3304) were cultured in complete EMEM (ATCC® 30-2003™) by adding 56 mL FBS (ATCC® 30-2020™) to a 500 mL bottle of the base medium. For BMEC (cAP-0001) and HUVEC (cAP-0002) cells were cultured in Endothelial Growth Medium (EGM, contains 10% serum and growth supplements, cAP-02). All cell culture and biological experiments were performed at 37°C in 5% CO2 in a cell culture incubator. For cell pretreatment, dZTP (Trilink, N-2003-1) was used.

##### Cell transfection

Cells were seeded at a density of 6–8 × $10^4$ cells per well in 48-well plates and allowed to attach and grow prior to experiments assessing immune activation in RAW264.7, RAW-Lucia ISG and HEK-Blue-mTLR7 cells. For mRNA translation assays using non-neuronal cells, a seeding density of 8-10 × $10^4$ cells per well in 24-well plates was used. Neural cells were seeded at ~70% confluence in 24-well plates for mRNA translation assays. Lipofectamine MessengerMAX was used as a standard transfection reagent for RNA and DNA transfection, following the manufacturer's protocol. For ribonucleoprotein (RNP) transfections, Lipofectamine CRISPRMAX was used according to the manufacturer's protocol. For RAW264.7, RAW-Lucia ISG and HEK-Blue-mTLR7 cells, the passage number did not exceed six.

##### Flow cytometry and fluorescence analysis

Cell culture medium was removed, and cells were detached using Trypsin-EDTA (Gibco, 25200056). Cells were analyzed directly using the BD Accuri™ C6 Flow Cytometer (BD Biosciences) to measure GFP fluorescence. Data was processed and analyzed using FlowJo software. Fluorescence imaging was performed using the KEYENCE BZX-810.

##### Preparing RNA by in *vitro* transcription

PCR DNA or linearized plasmids were used as templates for synthesizing mRNA through *in vitro* transcription (IVT). We utilized the HiScribe SP6 RNA Synthesis Kit (NEB, E2070) and set up 25 µL reactions, where were mixed thoroughly and incubated at 37°C for 2-4 hours. Following incubation, 25 µL of nuclease-free water and 2 µL of DNase I were added to each reaction to digest the DNA, followed by a 15 min incubation at 37°C. mRNA was purified using the Monarch RNA Cleanup Kit (NEB, T2040) and concentration was determined with a Thermo Scientific NanoDrop. For mRNA transcription, 0.4-0.8 µg of DNA were used per batch.

For small RNA (<200 nt transcription), 1 picomole of synthetic Ultramer™ DNA Oligonucleotides (IDT) was used as a template, with either the T7 High Yield RNA Synthesis Kit (NEB, E2040S) or the SP6 RNA Synthesis Kit used for RNA IVT.

Modified NTPs were used in place of standard NTPs to produce modified RNA with IVT. These included N1-methylpseudo-UTP (Jena Bioscience, NU-890), 2-thio-UTP (Jena Bioscience, NU-1151), pseudo-UTP (Jena Bioscience, NU-1139), 5-methoxy-UTP (Jena Bioscience, NU-972), GTPαS (Jena Bioscience, NU-409), CTPαS (Jena Bioscience, NU-436), N6-methyl-ATP (Apexbio, B7966) and ZTP (Trilink, N-1001; Jena Bioscience, NU-250). The quality of the mRNA was verified by gel electrophoresis.

5'-end Cap-0 synthesis was performed using Vaccinia Capping Enzyme (NEB #M2080). 25 µg of mRNA was diluted with nuclease-free water to a minimum volume of 15 µL, heated at 65°C for 5 min, and cooled on ice for 5 min. The denatured RNA was added to a 50 µL capping reaction following the manufacturer's protocol, incubated at 37°C for 30 min, and purified using the Monarch RNA Cleanup kit. The mRNA was eluted with 50 µL of nuclease-free water, and its concentration was determined with a NanoDrop. 3' Poly(A) tails were added to mRNA using the Poly(A) Polymerase Tailing Kit (LGC Biosearch Technologies, PAP5104H). A 50 µL reaction containing 12.5 µg of capped mRNA and 1.25 µL of RNase Inhibitor was incubated at 37°C for 30 min and purified using the Monarch RNA Cleanup Kit. The final mRNA was eluted with 50 µL of nuclease-free water, resulting in capped and tailed mRNA.

Purified mRNAs were stored at -80°C.

**Prepare dsRNA and gRNA**

For SpCas9 gRNA, 141.7 pmol of crRNA and 141.7 pmol of tracrRNA were mixed in RNase free water. The mixture was incubated at 70°C for 5 min and then allowed to cool to room temperature. For dsRNA preparation, 445 pmol of sense and antisense ssRNA were mixed in RNase free water and incubated at 70°C for 5 min and then allowed to cool to room temperature.

**Evaluate RIG-I activation by dsRNA**

The HEK293T cell line and its derivatives naturally express only a few pattern-recognition receptors (PRRs), including TLR3, TLR5, and NOD1. This characteristic makes them advantageous for studying specific signaling pathways upon transfection with genes encoding a given PRR. 80,000 HEK293T cells were plated in a 48-well plate. The following day, cells were transfected with 200 ng of IFNß-pGL3 (Addgene, 102597) and RIG-I (Addgene, 167289) plasmids using lipofection. 10 hours after the first transfection, dsRNA was lipofected into cells. After 24 hours, cells were washed with PBS, lysed, and 20 µL of the lysates were used to perform a Luciferase assay (Promega, E4030). Luminescence was measured using the Varioskan LUX Multimode Microplate Reader (ThermoFisher, US).

**Evaluate IFN responses in RAW-Lucia ISG cells.**

ISG cells were plated in 48-well plates. The following day, cells were transfected with various RNA constructs or treated with different doses of inhibitors, NTPs, dNTPs, RNA-induced PKR autophosphorylation inhibitor C16 (Sigma-Aldrich, 527450) and RIG-I inhibitor BX795 (Medchemexpress LLC, HY-10514). 10 hours after the initial lipofection, RNA was transfected using lipofection. At the designated time points post-treatment, culture media was collected to perform the Lucia luciferase assay (InvivoGen, rep-qlc4lg1). Luminescence was measured using either the Varioskan LUX Multimode Microplate Reader (ThermoFisher, US) or the BioTek Synergy H1 Hybrid Multi Mode Microplate Reader (BioTek).

**SEAP assay**

HEK-Blue TLR7 cells were plated at a density of $8 \times 10^4$ cells in a 48-well. Cells were treated with varying doses of IVT RNA or NTPs. After the designated incubation time, 50 µL of culture media was collected and transferred to a new 96-well plate. Next, 100 µL Quanti-Blue (InvivoGen, rep-qbs) was added to each well containing the media, and the plate was incubated at 37°C until a visible color change occurred. Absorbance was measured using a microplate reader at 620-650 nm.

**Statistical Analysis and Graphical Illustrations.**

Curve plotting and statistical analyses were performed using Prism 8 (GraphPad, La Jolla, CA). Graphical illustrations were created using PyMOL (DeLano Scientific LLC), Microsoft PowerPoint, and ChemDraw (PerkinElmer).

**Figures**

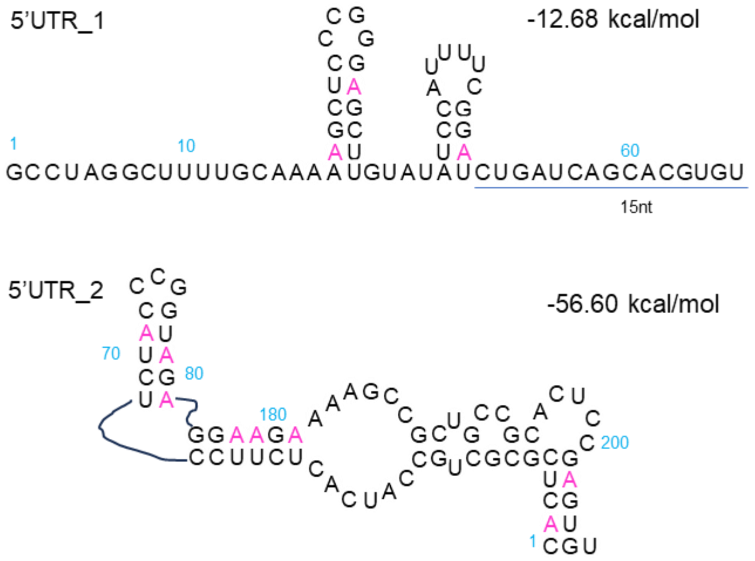


**Fig. s1 Structural analysis of UTR sequence.** Secondary structures were predicted using RNAfold web server. The thermodynamic free energy (in kcal/mol) represents the ensemble free energy at default NaCl concentrations (~1 M). Calculations were performed at 37℃ using standard energy parameters.

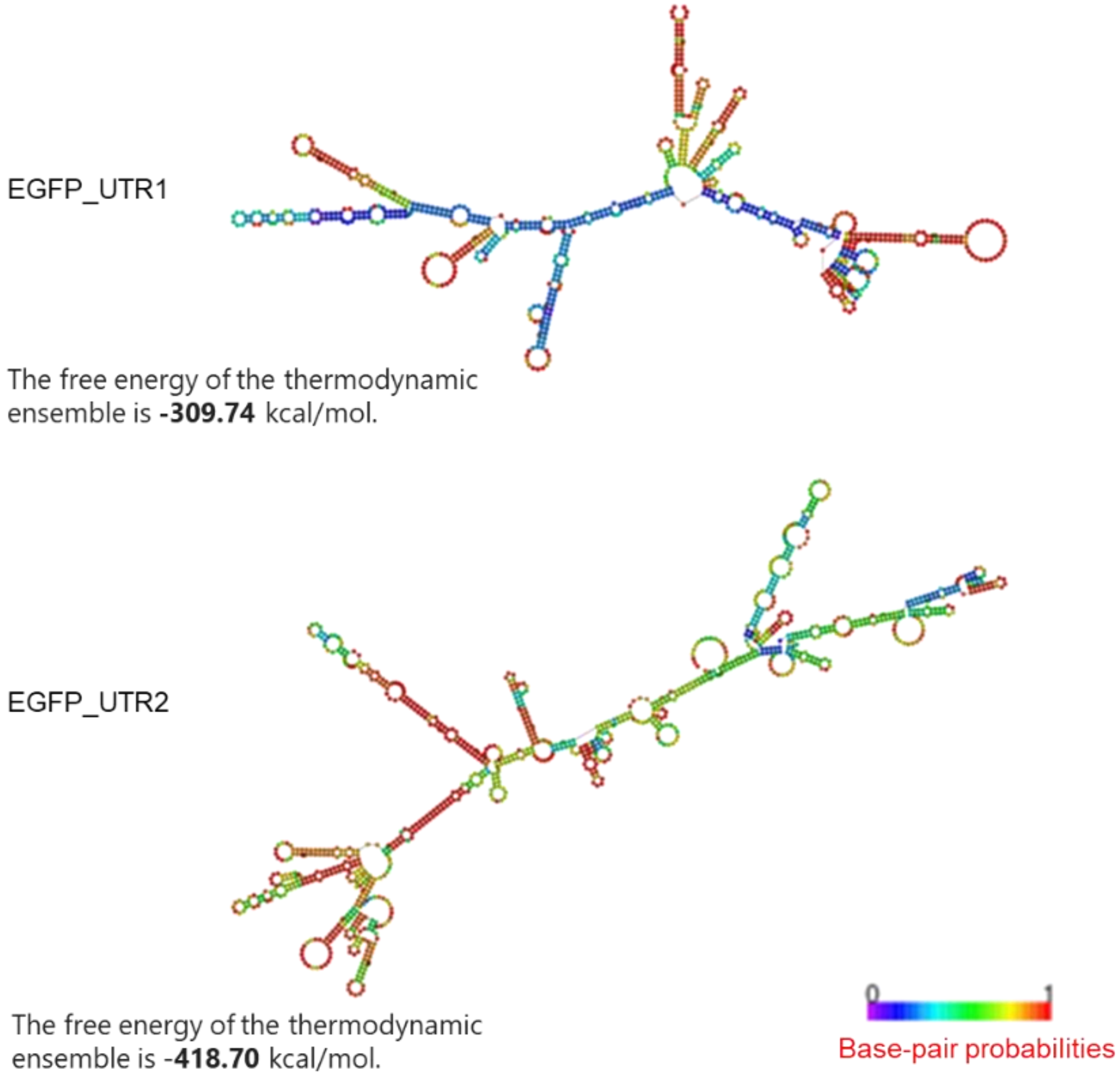


**Fig. s2 Structural analysis of EGFP mRNA with UTR1 and UTR2 sequences.** Secondary structures were predicted using the RNAfold web server. Energy parameters (in kcal/mol) were calculated at a default NaCl concentration of ~1 M and measured at 37°C.

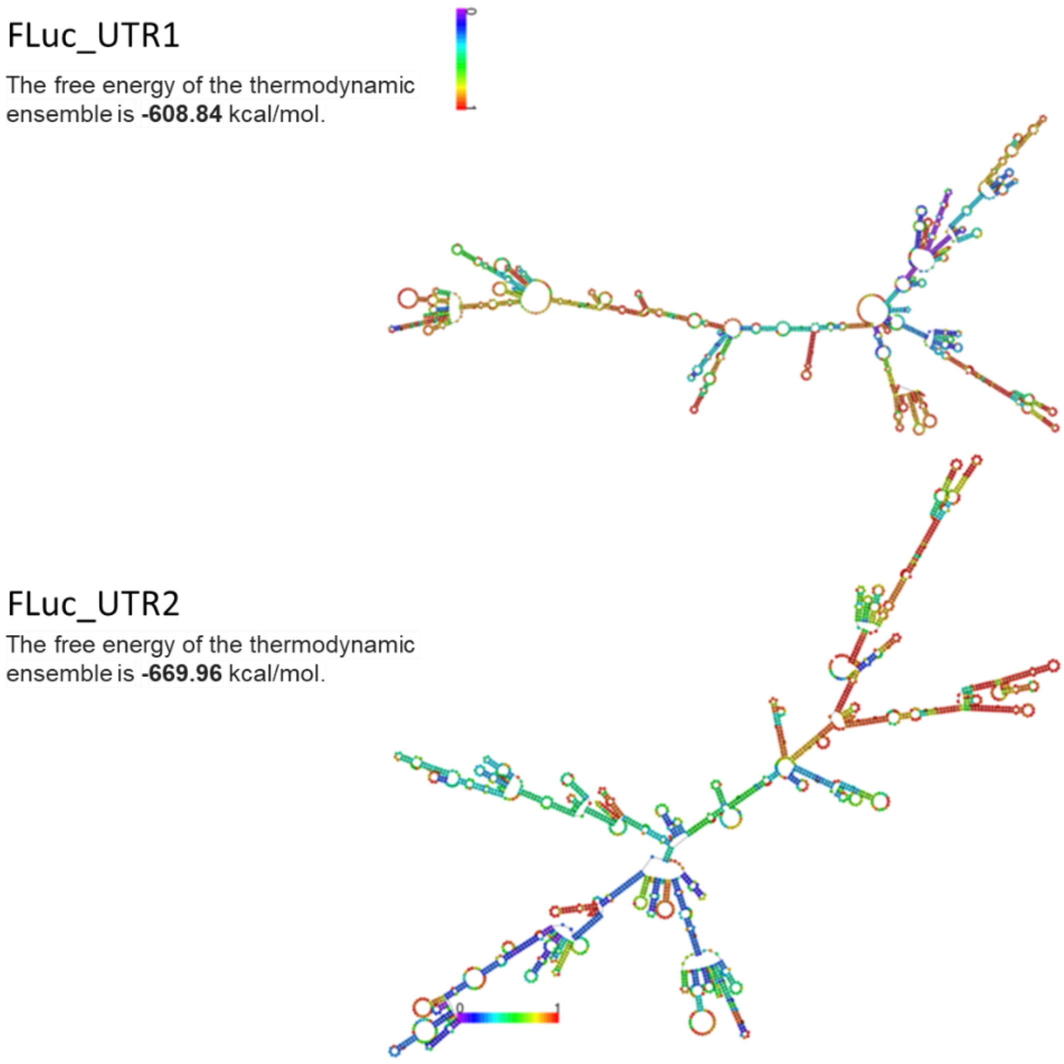


**Fig. s3 Structural analysis of FLuc mRNA with UTR1 and UTR2 sequences.** Secondary structures were predicted using the RNAfold web server. Energy parameters (in kcal/mol) were calculated at a default NaCl concentration of ~1 M and measured at 37°C.

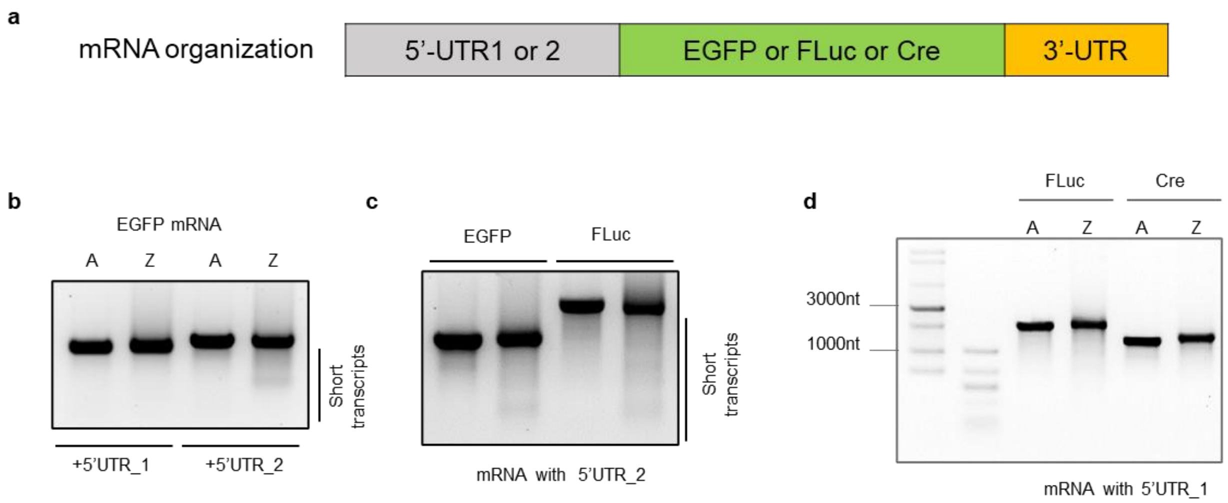


**Fig. s4 Influence of UTR sequences on IVT transcripts. (a)** Schematic representation of the mRNA constructs used. (**b)** Gel analysis of EGFP mRNA with different UTR sequences. (**c)** Gel analysis of EGFP mRNA and FLuc mRNA with the UTR2 sequence. (**d)** Gel analysis of FLuc mRNA and Cre mRNA with the UTR1 sequence.

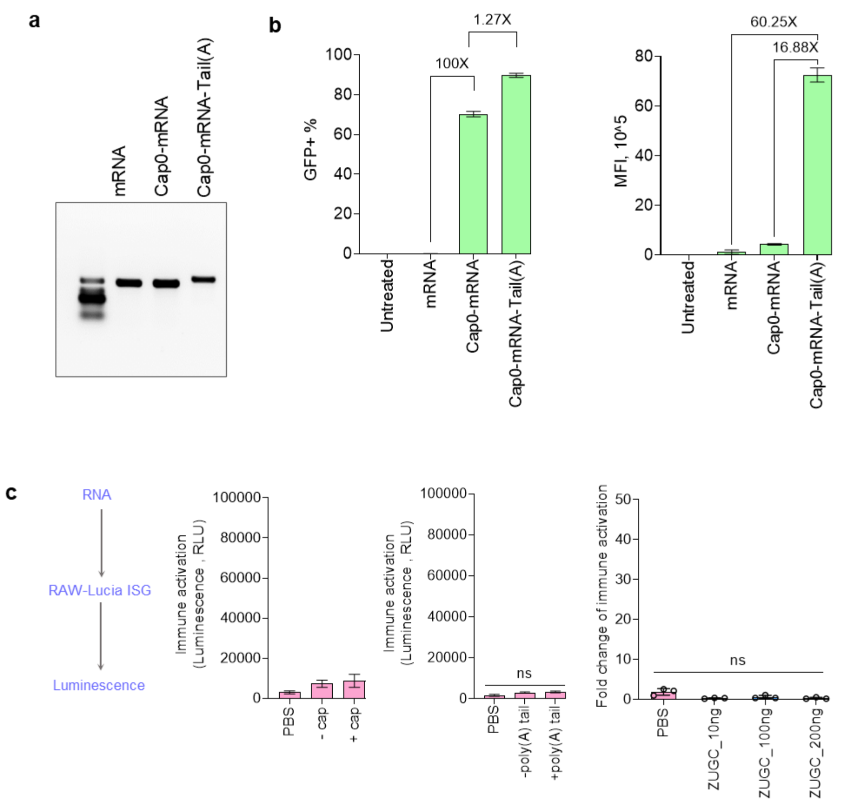


**Fig. s5 Influence of capping and tailing on EGFP Z-mRNA properties.** (**a)** Gel analysis of Z-mRNA with or without capping and tailing. **(b)** Expression analysis of EGFP Z-mRNA using HEK293T cells following lipofection (n = 3). (**c)** Immune activation mediated by various Z-mRNA constructs (n = 3). Statistical analysis was performed using an ordinary one-way ANOVA with Sidak's multiple comparisons test. *P* values of <0.05 were considered statistically significant. ns, not significant. All data is presented as mean ± SD.

RAW264.7 transfected with EGFP mRNA

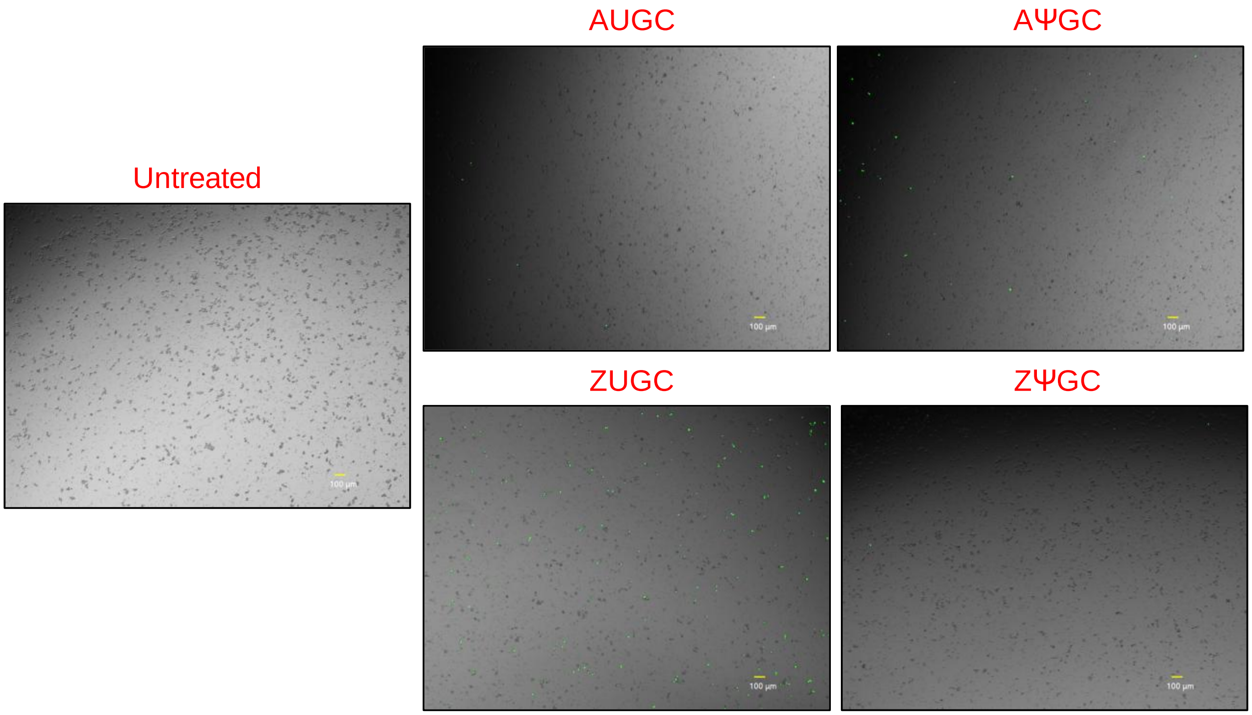


**Fig. s6 Representative merged brightfield and GFP images of RAW264.7 cells transfected with various mRNA constructs.**

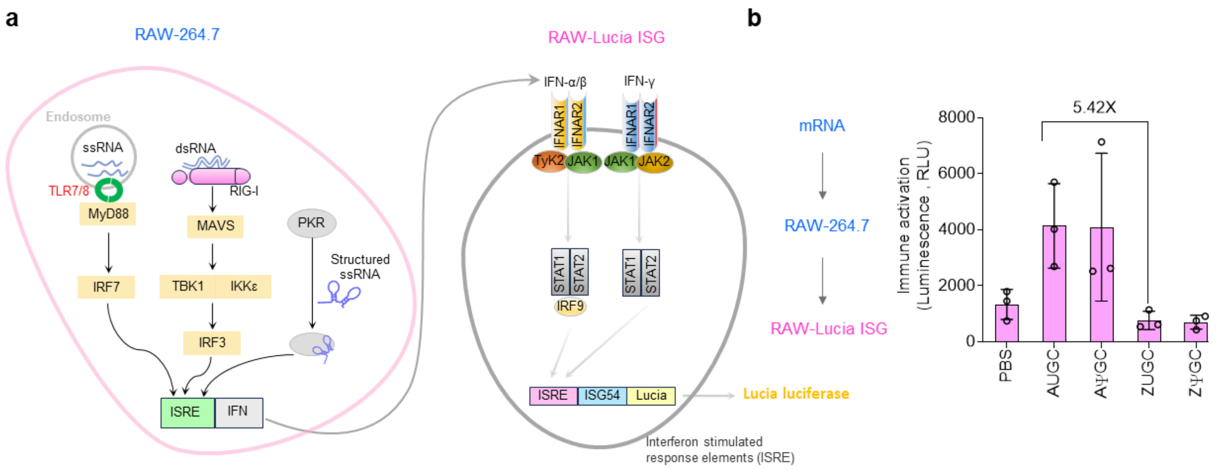


**Fig. s7 Immune activation mediated by various IVT mRNA in the macrophage RAW264.7 cell line.** (**a**) Schematic representation of the sensing pathway for exogenous RNA in RAW264.7 cells and the IFN sensing pathway in RAW-Lucia ISG cells. (**b**) Immune activation mediated by various IVT mRNA constructs in RAW264.7 (n = 3).

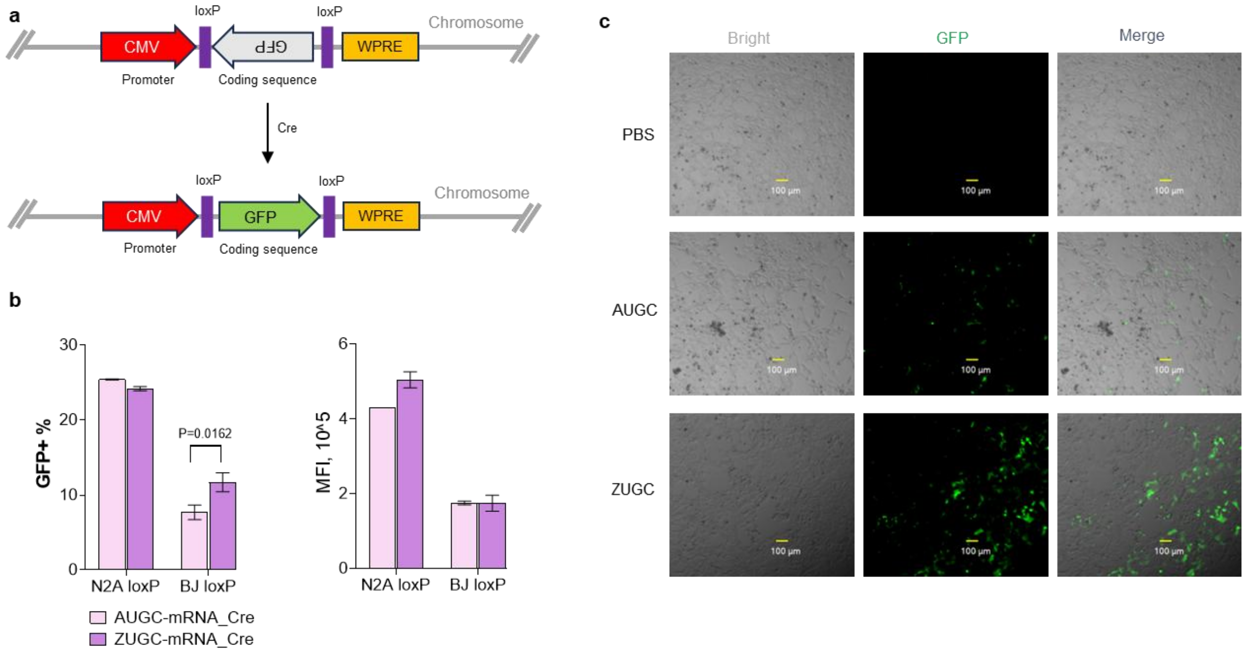


**Fig. s8 Cre mRNA activity in EGFP-loxP reporter cell lines.** (**a**) Schematic representation of Cre recombinase-mediated inversion of the EGFP cassette to generate active EGFP expression. (**b**) Activation efficiency and mean of fluorescence intensity (MFI) of GFP+ cells mediated by various Cre-mRNA constructs in N2A and BJ reporter cell lines. Data are presented as mean ± SD (n = 3). *P* values were analyzed using 2-way ANOVA Dunnett's multiple comparisons test. *P* values of $<0.05$ were considered statistically significant. (**c**) Representative brightfield, GFP, and merged images of BJ reporter cells treated by various Cre-mRNA constructs.

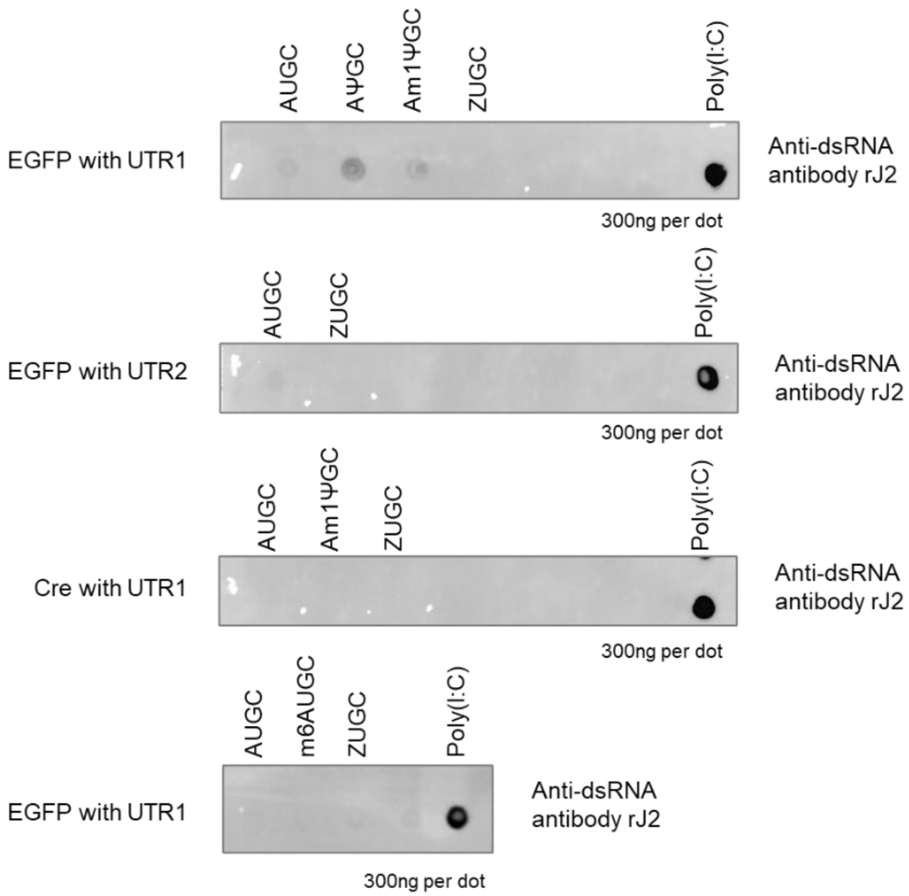


**Fig. s9 Detection of dsRNA using a dot blot assay.** Dot blot analysis was performed with an anti-dsRNA monoclonal antibody (rJ2).

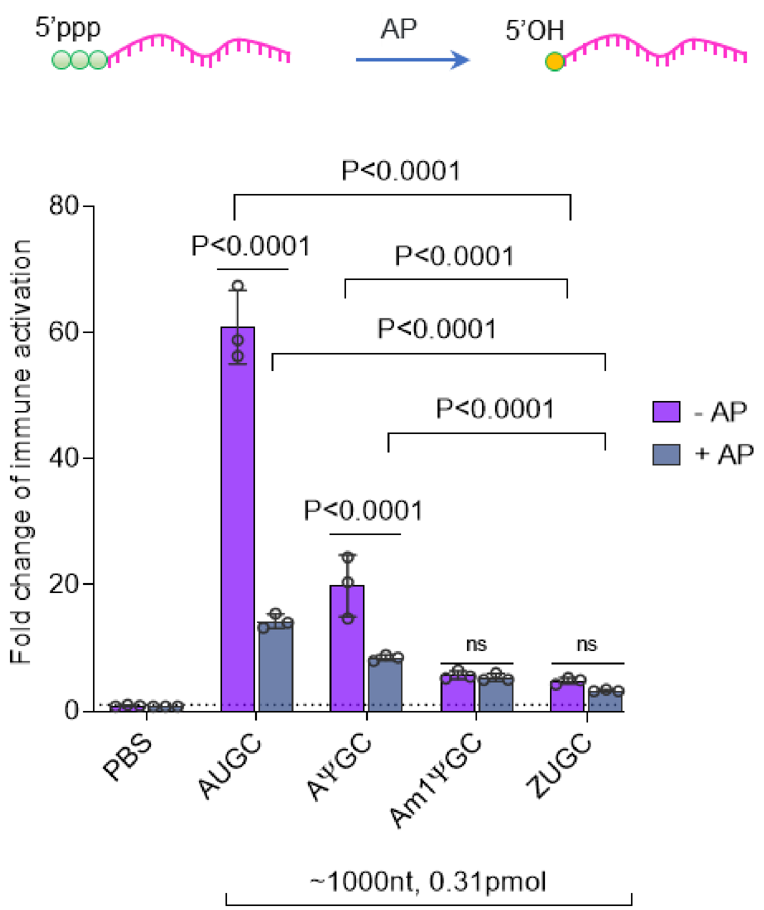


**Fig. s10 Immune activation induced by various mRNA constructs with or without Antarctic phosphatase (AnP) treatment.** Non-capped mRNA was used in this analysis (n = 3). Data are presented as mean ± SD. Statistical analysis was performed using a 2-way ANOVA with Sidak's multiple comparisons test. *P* values of <0.05 were considered statistically significant. ns, not significant.

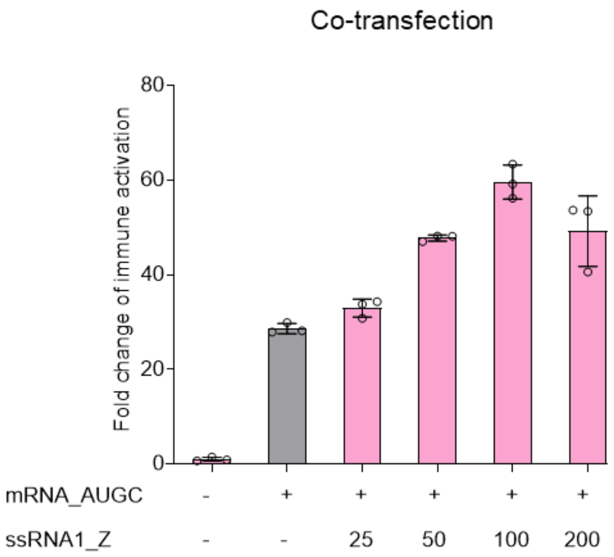


**Fig. s11 Immune activation induced by co-transfection of AUGC RNA and ZUGC RNA in ISG cells.** "+" indicates 100 ng of RNA treatment. EGFP AUGC-mRNA and ZUGC-ssRNA1 were used in this experiment. Data are presented as mean ± SD (n = 3).

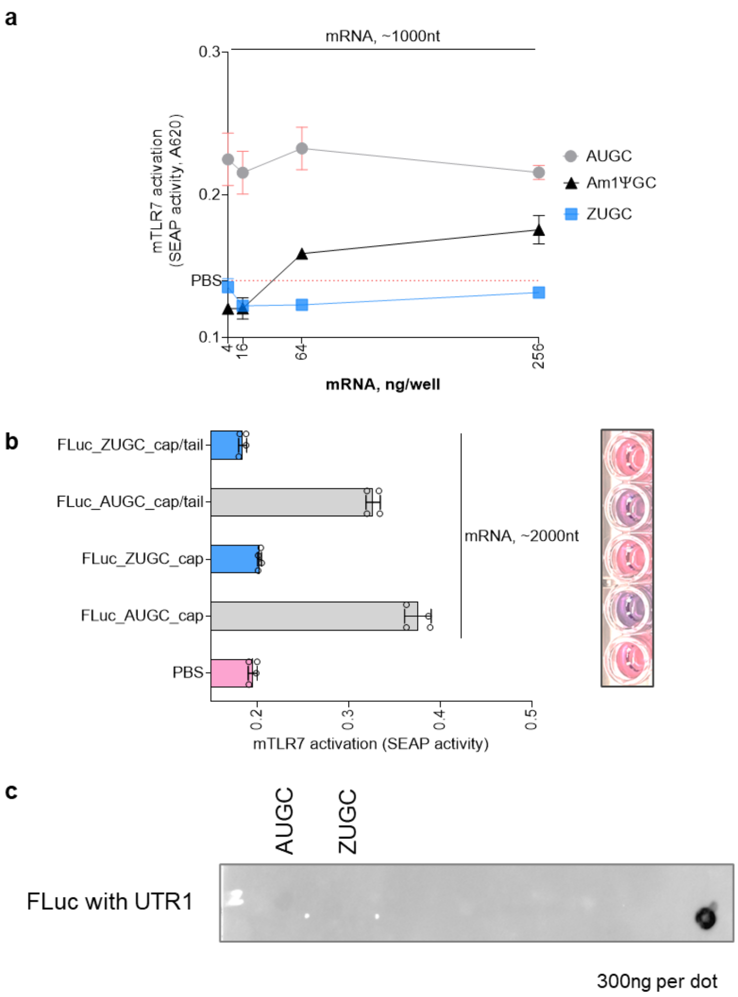


**Fig. s12 mTLR7 activation in HEK-Blue-mTLR7 cells.** (**a**) mTLR7 activation induced by various EGFP mRNA constructs (n = 3). (**b**) mTLR7 activation induced by various FLuc mRNA constructs (n = 3). (**c**) dsRNA analysis of FLuc mRNA. All data is presented as mean ± SD.

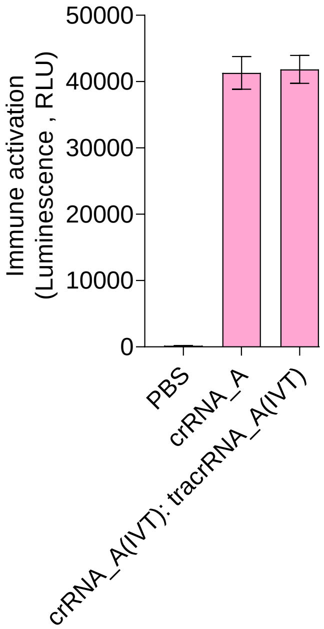


**Fig. s13 Immune activation induced by crRNA or gRNA.** gRNA is formed by hybridizing crRNA_A with tracrRNA_A. Data are presented as mean ± SD (n = 3).

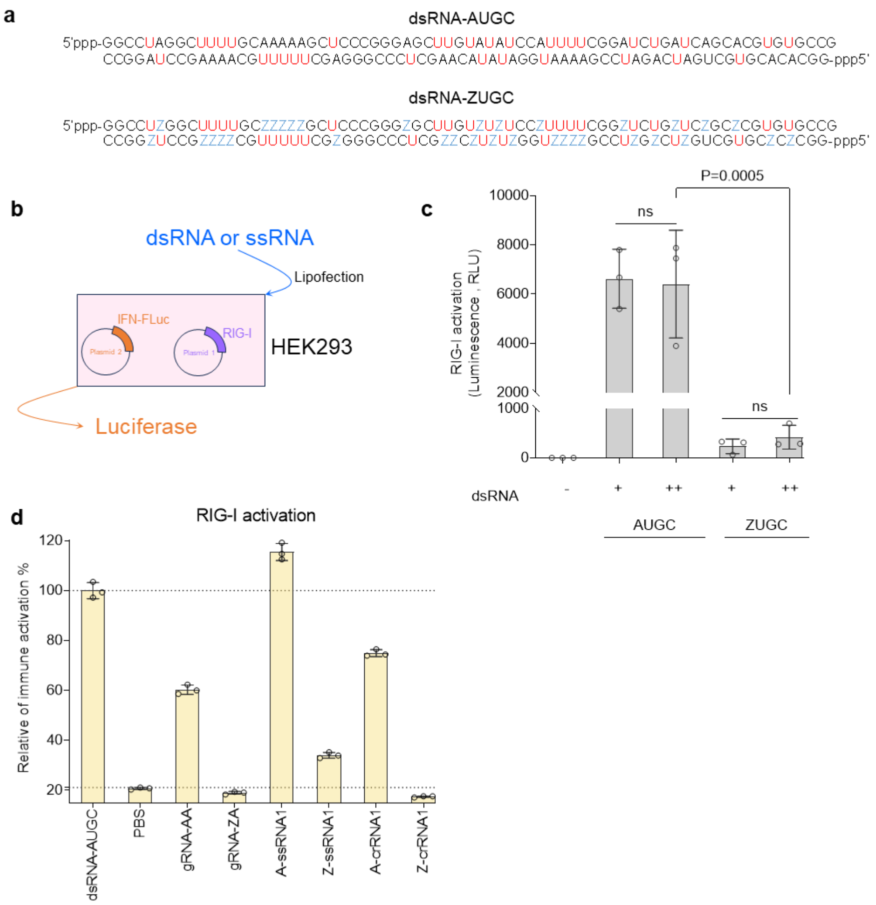


**Fig. s14 RIG-I activation mediated by AUGC or ZUGC ssRNA and dsRNA. (a)** dsRNA sequences used for RIG-I activation. **(b)** Design of RIG-I activation assay. **(c)** RIG-I activation induced by dsRNA. "+" indicates 100 ng dsRNA was used per well. (**d**) ssRNA also can evade RIG-I sensing to reduce immune response. Data are presented as mean ± SD (n = 3). For (c), statistical analysis was performed using an ordinary one-way ANOVA with Tukey's multiple comparisons test. *P* values of <0.05 were considered statistically significant. ns, not significant.

**a**

sgRNA_SYN (Genscript)
5'-
mG*mU*mA*rArArCrArArArGrCrArUrArGrArCrUrGrAr
GrUrUrUrUrArGrArGrCrUrArGrArArArUrArGrCrArArGr
UrUrArArArArUrArArGrGrCrUrArGrUrCrCrGrUrUrArUr
CrArArCrUrUrGrArArArArArGrUrGrGrCrArCrCrGrArGr
UrCrGrGrUrGrCrU*mU*mU*mU-3'

tracrRNA_SYN (IDT, Alt-R$^{TM}$)
5'-
rArGrCrArUrArGrCrArArGrUrUrArArArArUrArArGrGrCrU
rArGrUrCrCrGrUrUrArUrCrArArCrUrUrGrArArArArArGrU
rGrGrCrArCrCrGrArGrUrCrGrGrUrGrCrUrUrU-3'

Note: IDT stated that the modifications on this tracrRNA are proprietary and cannot be shared.

**b**

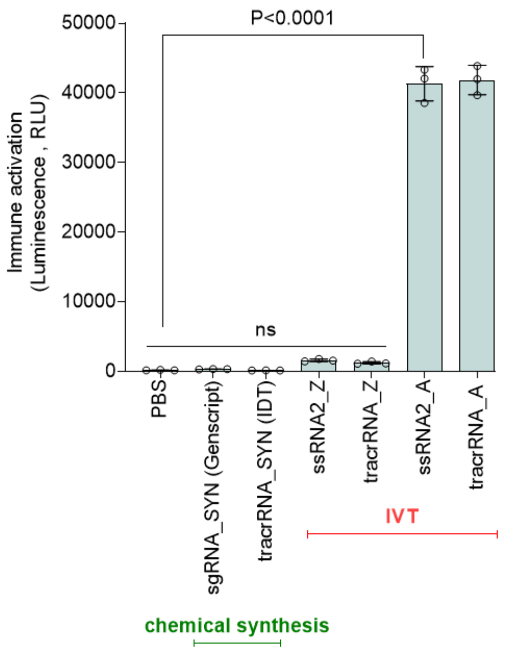


**Fig. s15 Comparison of IVT Z-RNA with two chemically synthesized RNAs in ISG cells.** (a) Sequences of tracrRNAs ordered from Genscript and IDT. (b) Immune activation induced by chemical synthezied or IVT RNA. Data are presented as mean ± SD (n = 3). Statistical analysis was performed using an ordinary one-way ANOVA with Sidak's multiple comparisons test. *P* values of <0.05 were considered statistically significant. ns, not significant.

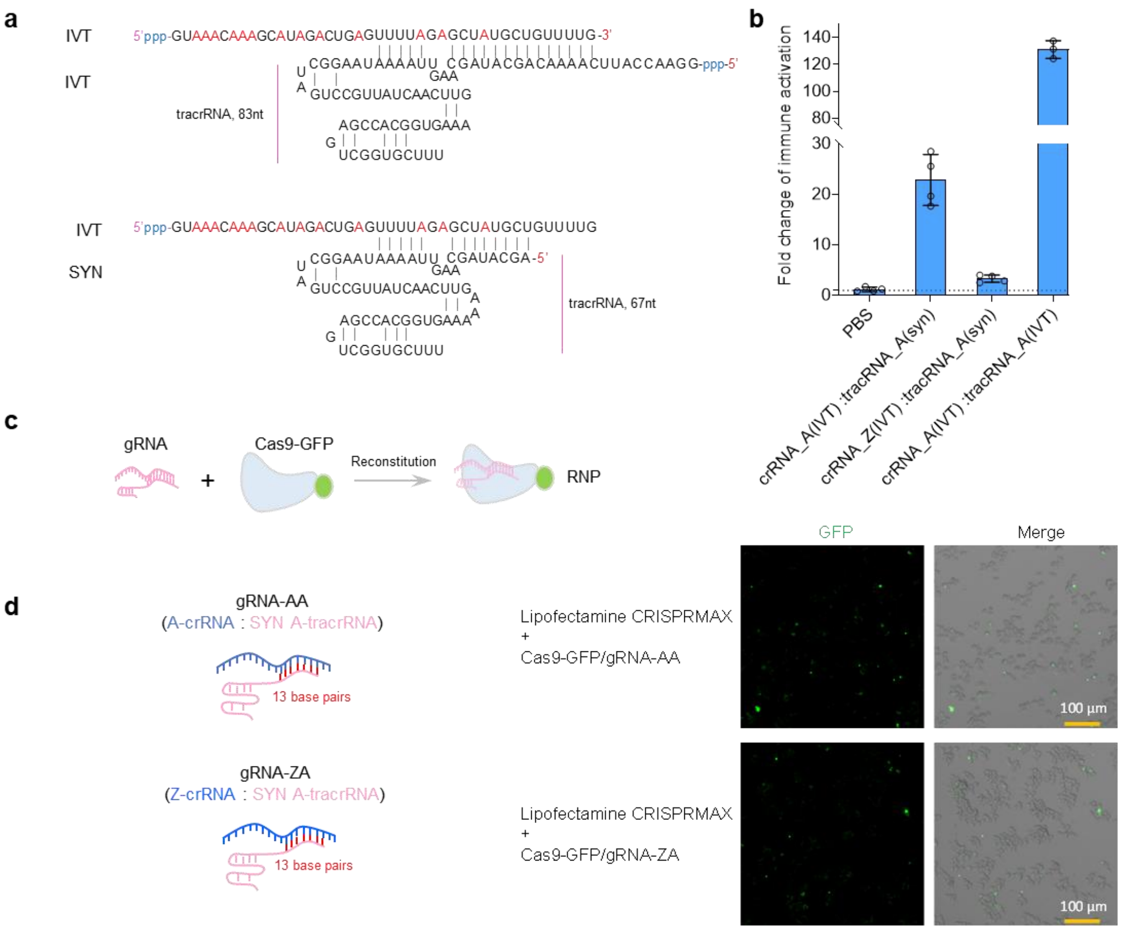


**Fig. s16 Immune activation induced by small dsRNA with or without Z incorporation.** (**a**) Schematic representation of gRNA generated by annealing crRNA and tracrRNA. IVT, in vitro transcription; SYN, chemical synthesis. (**b**) Immune activation induced by different gRNAs was tested in ISG cells. Data are presented as mean ± SD (n = 4). (**c**) Ribonucleoprotein (RNP) complexes were prepared by reconstituting gRNA and Cas9-GFP protein. (**d**) Representative GFP and merged brightfield and GFP images of ISG cells treated with RNPs.

ssRNA_1

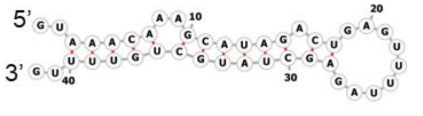

ssRNA_2

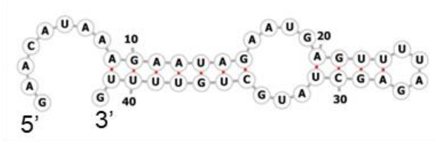

tracrRNA_SpCas9

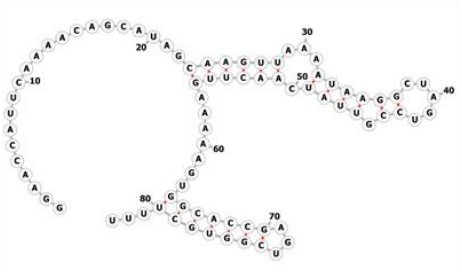

**Fig. s17 Predicted secondary structures of crRNA and tracrRNA used in Fig. 3a.** Secondary structures were predicted using the RNAfold web server. Parameters were calculated at a default NaCl concentration of ~1 M and measured at 37℃.

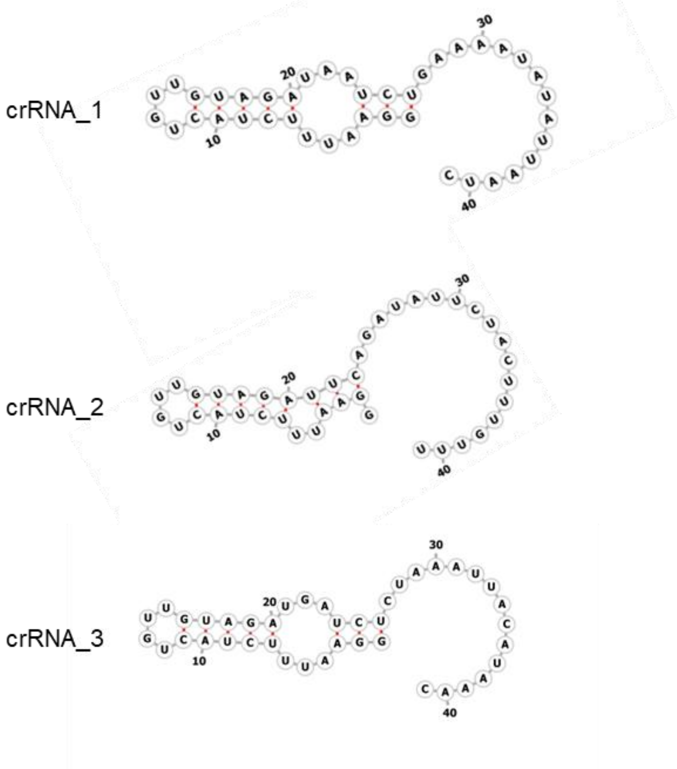


**Fig. s18 Predicted secondary structure of crRNA used in Fig. 4a.** Secondary structures were predicted using the RNAfold web server.

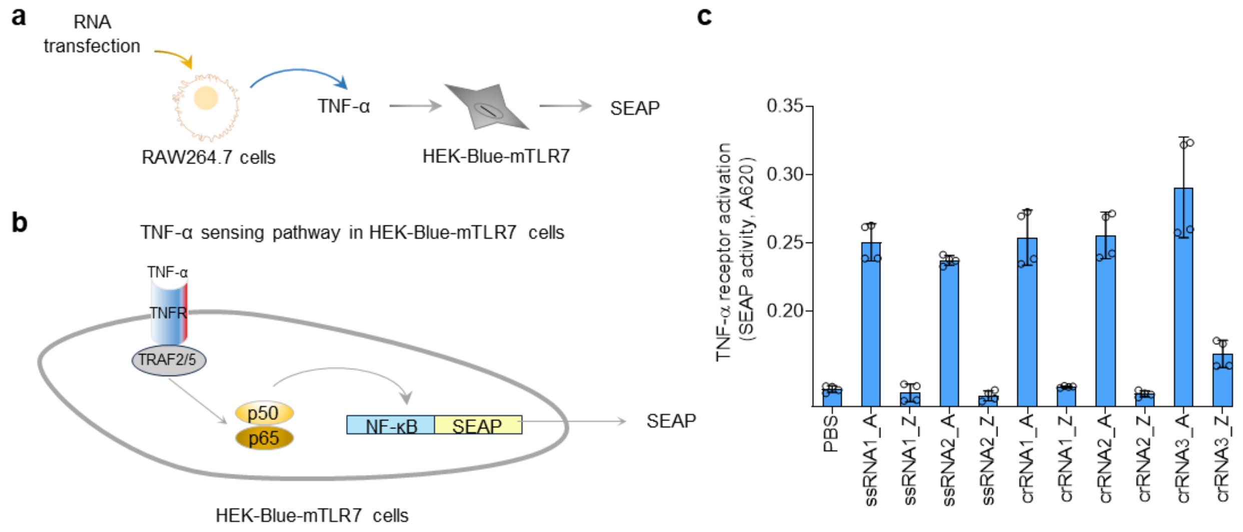


**Fig. s19 TNF-α cytokine production induced by AUGC or ZUGC RNA in RAW264.7 cells.** **(a)** TNF-α production was measured using the SEAP assay. (**b**) Schematic illustration of TNF-α sensing pathway in HEK-Blue-mTLR7 cells. (**c**) activation of TNF-α receptor was indicated by SEAP activities. Data are presented as mean ± SD (n = 4).

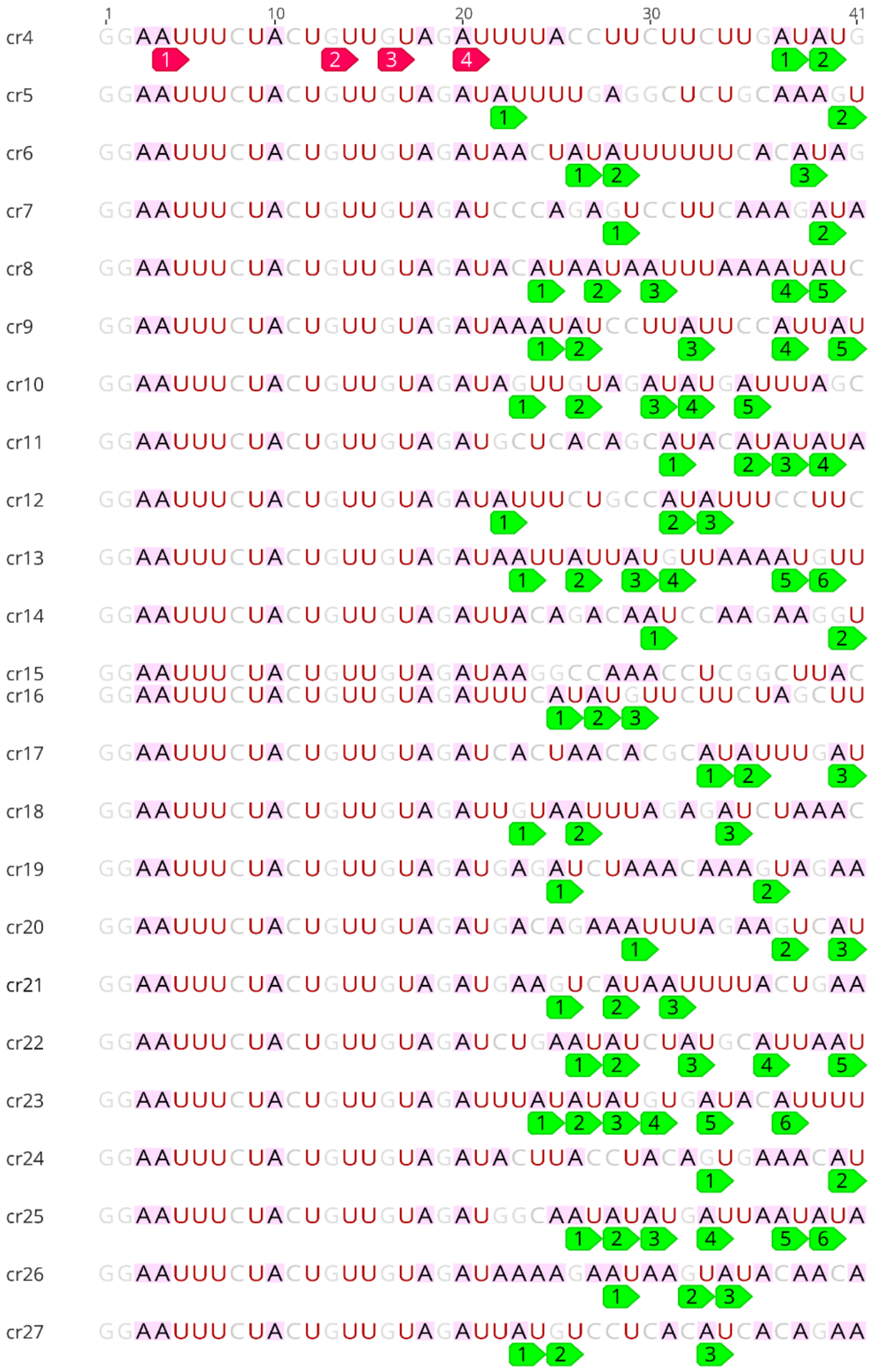

**Fig. s20 24 crRNA sequences used in Fig. 4e.** U is highlighted in red; A is highlighted in black with a pink background; numbered red frames indicate RU motifs in the fixed region; numbered green frames indicate RU motifs in the variable region.

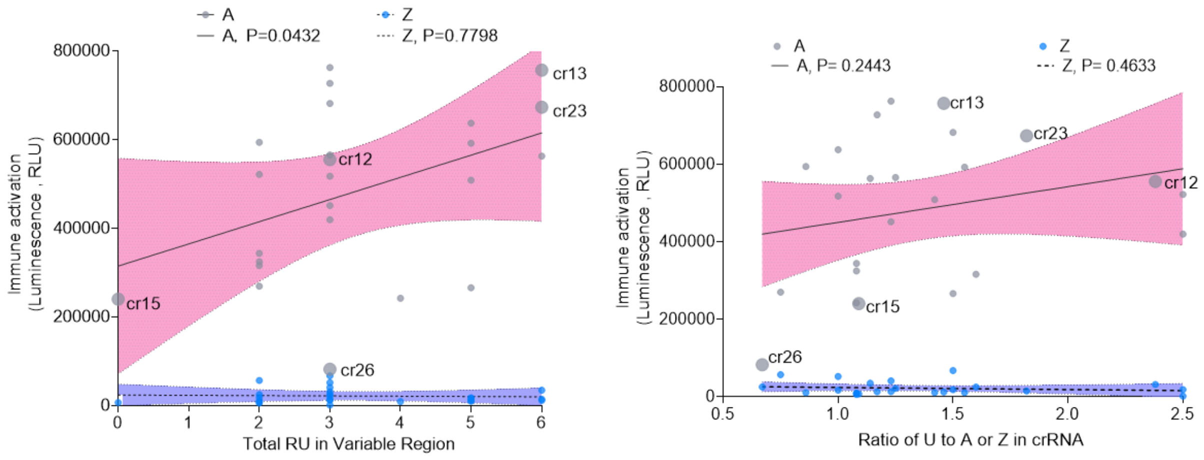


**Fig. s21 Correlation between immune activation and RU content in the variable regions of ssRNA used in Fig. 4f.** Linear regression analysis was used to analyze data. *P* values of <0.05 were considered statistically significant.

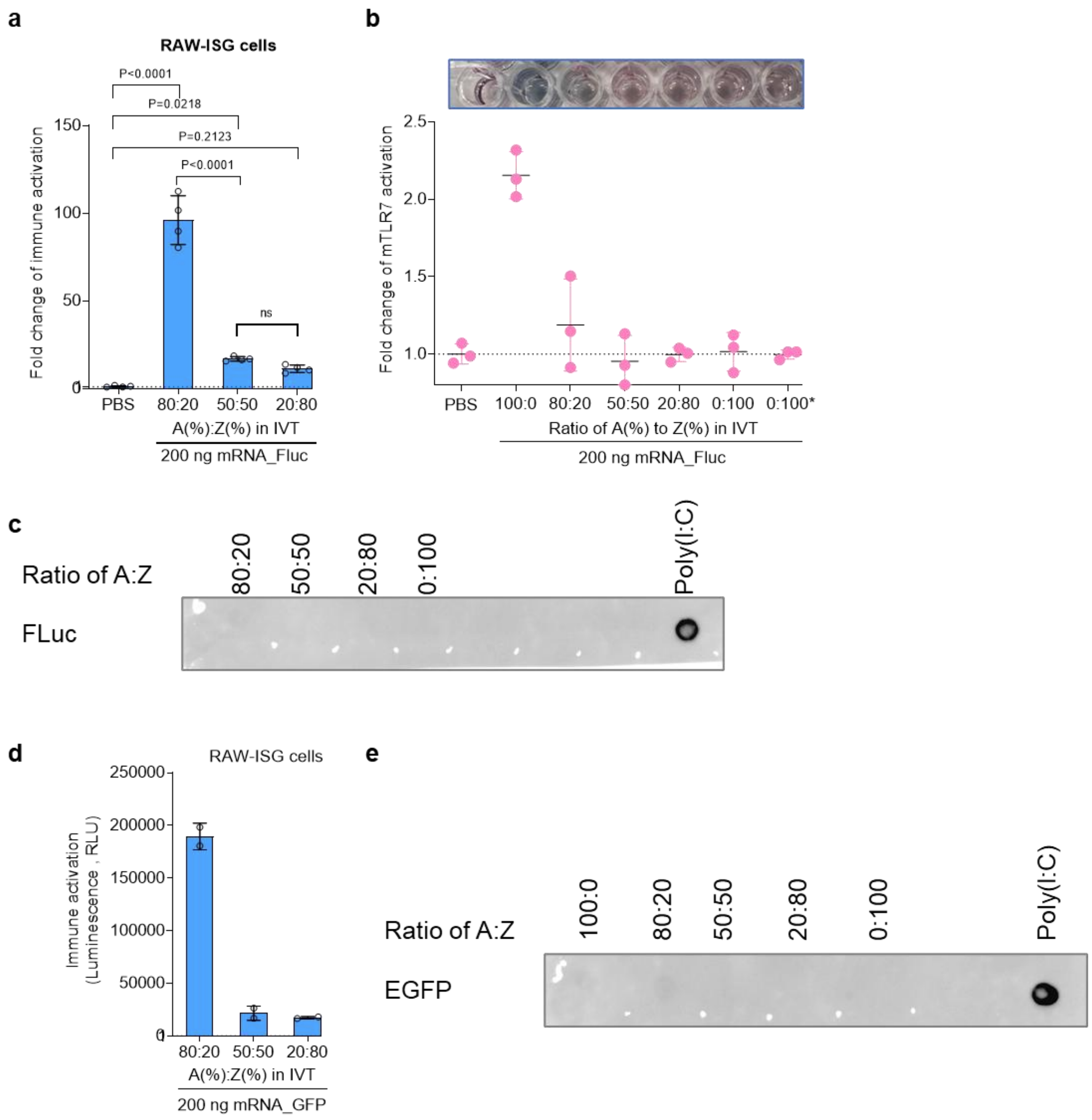


**Fig. s22 Evaluating non-complete substitution of Z in IVT. (a)** Immune activation in ISG cells mediated by FLuc mRNA with varying Z substitution percentages (n = 4). Statistical analysis was performed using ordinary one-way ANOVA with Tukey's multiple comparisons test. *P* values of <0.05 were considered statistically significant. ns, not significant. (**b**) mTLR7 activation mediated by FLuc mRNA with varying Z substitution percentages (n = 3). (**c**) dsRNA detection in FLuc mRNA used in (b). (**d**) Immune activation in ISG cells mediated by EGFP mRNA with varying Z substitution percentages (n = 2). (**e**) dsRNA detection in FLuc mRNA used in (d). All data is presented as mean ± SD.

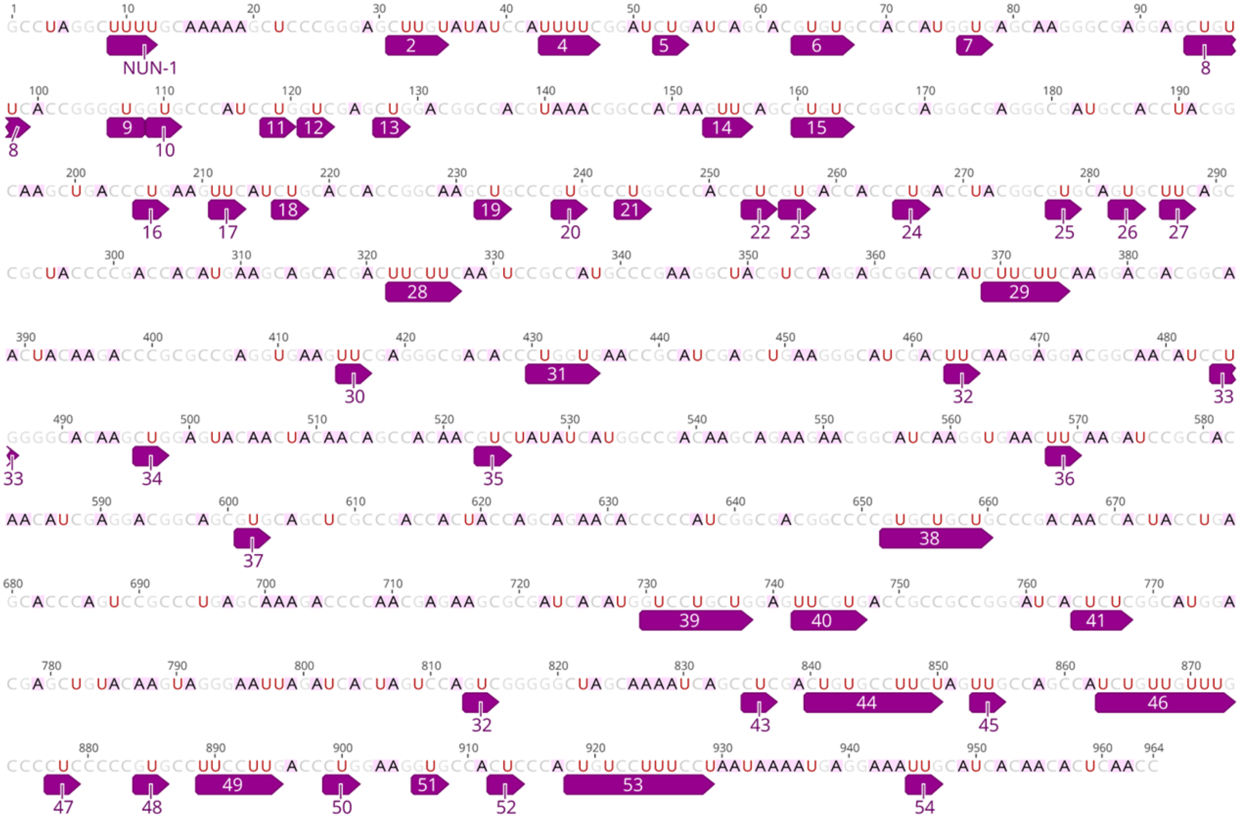

**Fig. s23 Schematic illustration of NUN motifs in EGFP-UTR1 sequences used in Fig. 2.** This sequence contains 227 As, 167 Us, 42 GU motifs, 34 AU motifs, 64 GG motifs, 75 AC motifs, 16 UUC motifs, 6 UUU motifs, and 13 CUU motifs. Numbered purple frames indicate motifs that can potentially generate NUN nucleotides.

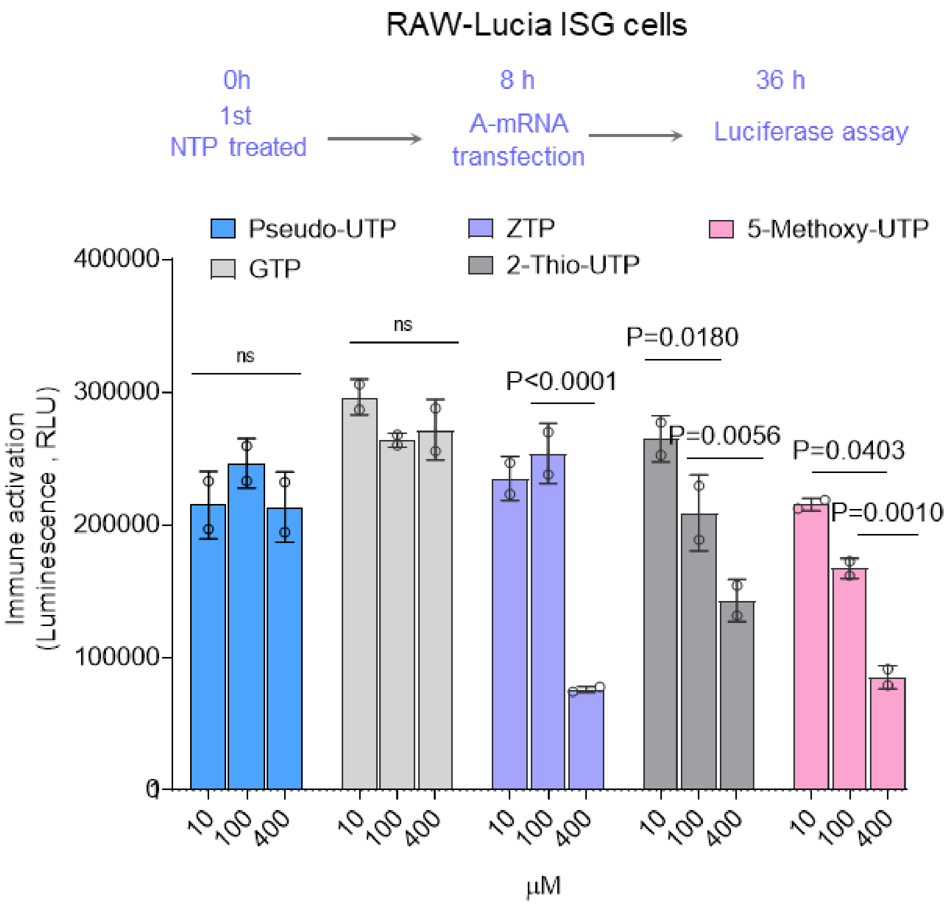


**Fig. s24 Screening of NTPs for inhibitory effects on immune activation induced by lipofected AUGC-ssRNA.** NTPs were directly added to the culture medium. 100 ng of mRNA was used for each treatment. Data are presented as mean ± SD (n = 2). Statistical analysis was performed using 2-way ANOVA with Tukey's multiple comparisons test. *P* values of <0.05 were considered statistically significant. ns, not significant. Cre mRNA was used in this evaluation.

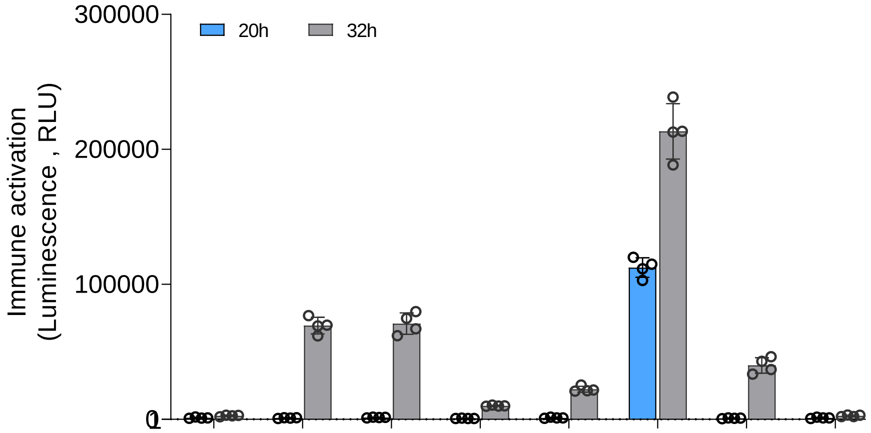


| | | | | | | | | | |
|---|---|---|---|---|---|---|---|---|---|
| 1st | PBS | + | + | - | - | - | - | + | + |
| | Z-mRNA | - | - | + | - | - | - | - | - |
| | ZTP | - | - | - | + | - | - | - | |
| | dZTP | - | - | - | - | + | - | - | - |
| | A-mRNA | - | - | - | - | - | + | - | - |
| | Lipofection | - | + | + | + | + | + | + | - |
| 2nd | PBS | + | - | - | - | - | - | - | + |
| | A-mRNA | - | + | + | + | + | + | + | - |
| | Lipofection | + | + | + | + | + | + | + | - |

**Fig. s25 Original data used in Fig. 5b.** The chart illustrates experimental conditions comparing the effects of Z-mRNA, ZTP, dZTP, and A-mRNA treatments in the presence or absence of lipofection. The x-axis indicates experimental groups, while the y-axis shows the measured outcome. The conditions are divided into two stages (1st and 2nd). 1st Stage: Positive (+) or negative (-) signs represent the presence or absence of specific treatments, including PBS, Z-mRNA, ZTP, dZTP, A-mRNA, and lipofection. 2nd Stage: Only PBS and A-mRNA with or without lipofection were included. Data are presented as mean ± SD (n = 4). Cre mRNA was used in this evaluation.

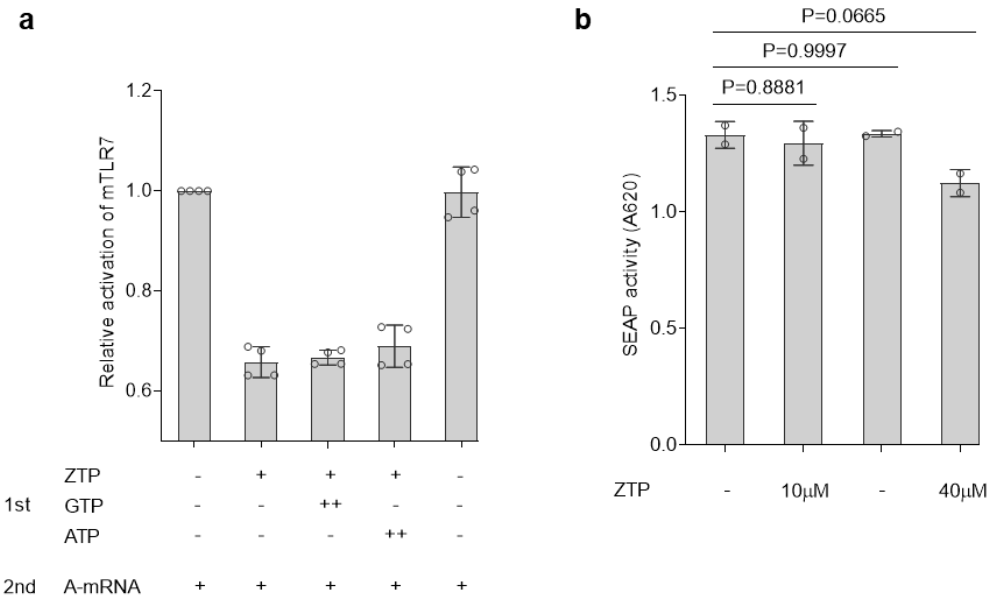


**Fig. s26 Effects of NTPs on immune activation. (a**) Influence of GTP and ATP on ZTP-mediated inhibition of immune activation (n = 4). For NTPs, "+" indicates a dose 4 μM. For A-mRNA, "+" indicates 50 ng EGFP mRNA was used (~156fmol). The conditions are divided into two stages (1st and 2nd). Positive (+) signs represent the presence or dose of treatment. Negative (-) signs represent absence of specific treatments. (**b**) Influence of ZTP on positive SEAP reaction control (n = 2). All data is presented as mean ± SD. For (b), statistical analysis was performed using an ordinary one-way ANOVA with Dunnett's multiple comparisons test. *P* values of <0.05 were considered statistically significant.

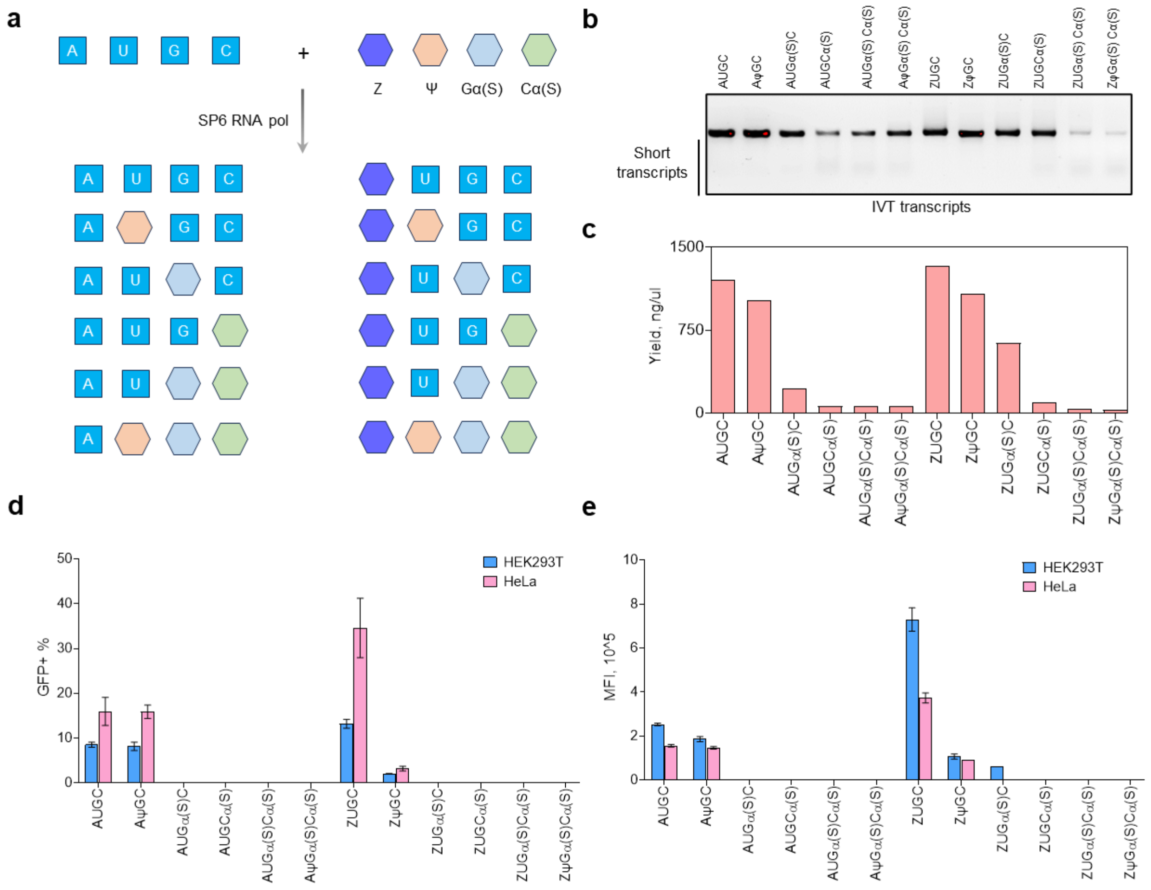


**Fig. s27 Screening mRNA with multiple modifications. (a)** Design of RNA transcription using various NTPs, including Gα(S) [GTP α(S)] and Cα(S) [CTPα(S)]. (**b**) Gel analysis of IVT products from (a). (**c**) Transcription yield of quantified from (b). (**d, e**) mRNA expression analysis after lipofection into HEK293T and HeLa cells (n = 3). Data are presented as mean ± SD.

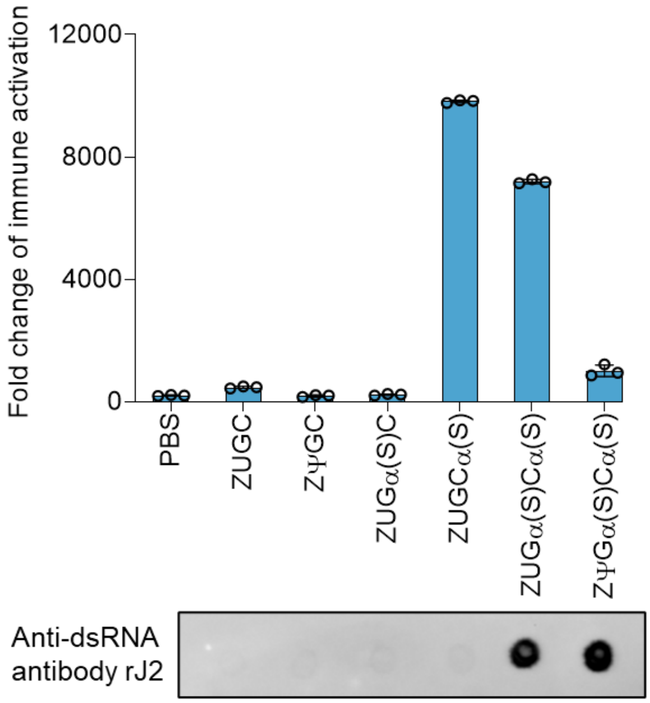


**Fig. s28 Immune activation in ISG cells transfected with various Z-mRNAs generated in (Fig. s27b).** Data are presented as mean ± SD (n = 3).

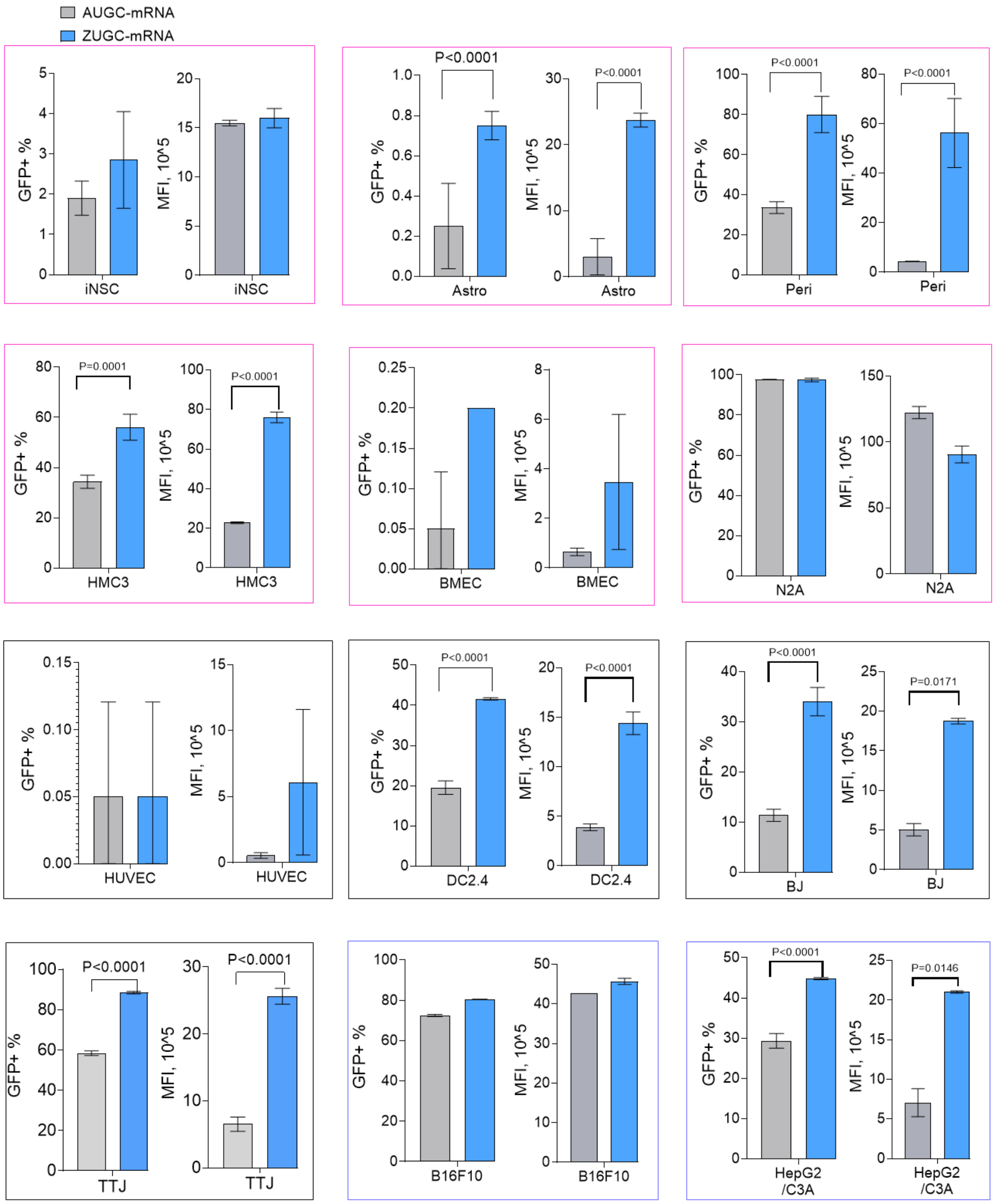


**Fig. s29 EGFP A-mRNA and Z-mRNA expression in various cell lines.** Data are presented as mean ± SD (n = 3). Statistical analysis was performed using ordinary one-way ANOVA with Sidak's multiple comparisons test. *P* values of <0.05 were considered statistically significant.

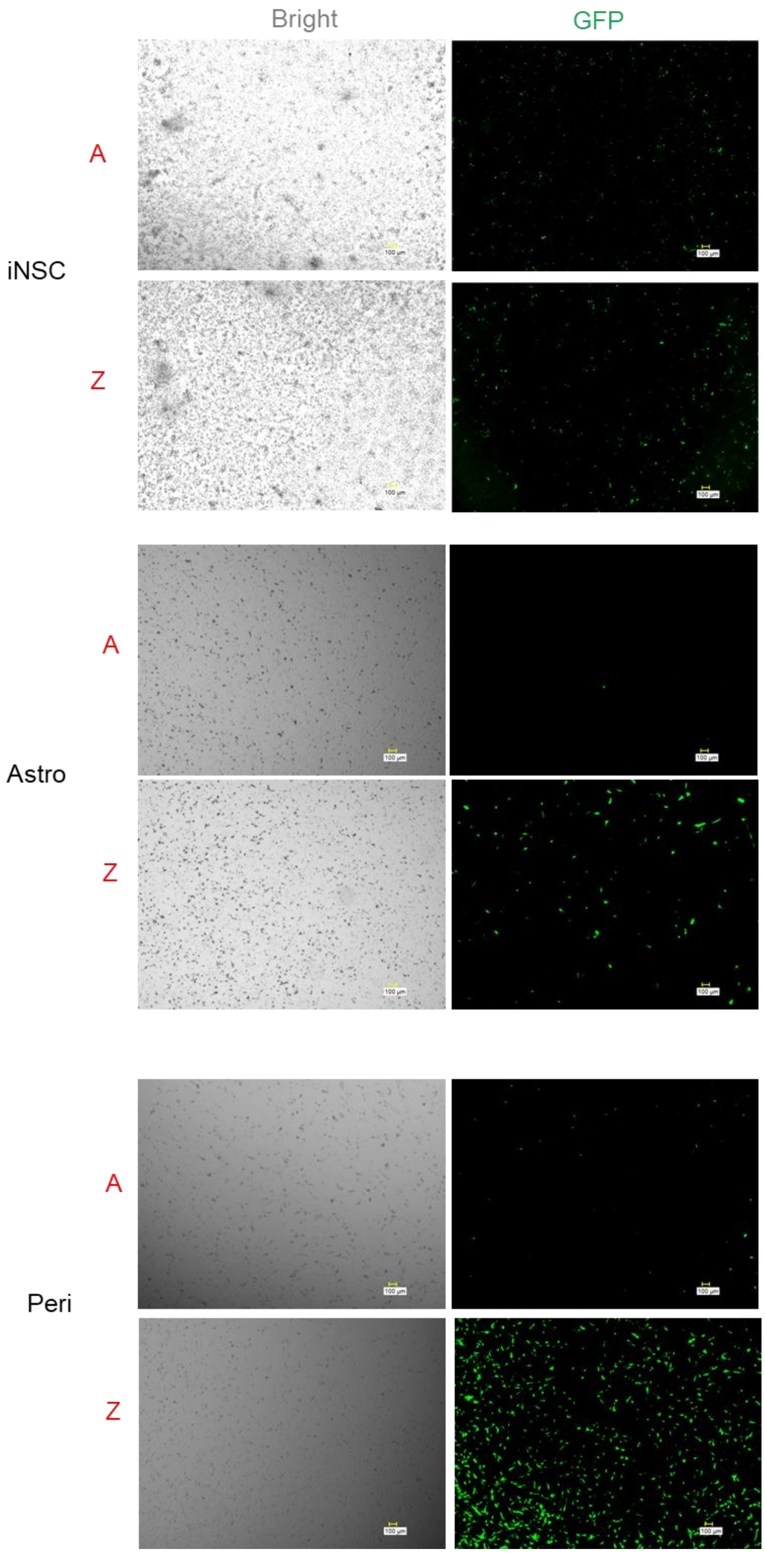
Bright
GFP
iNSC
A
Z
Astro
A
Z
Peri
A
Z

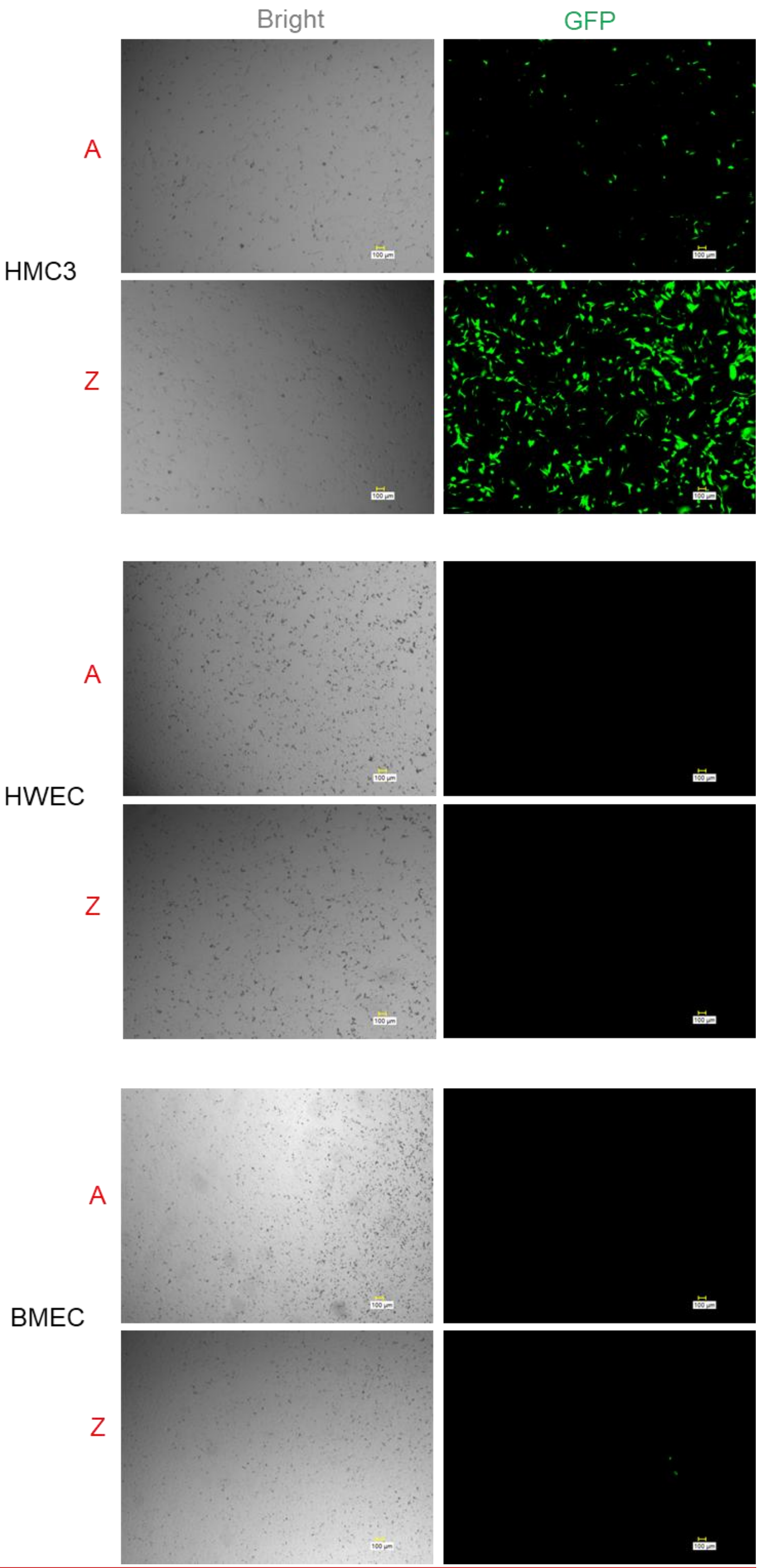
Bright
GFP
A
HMC3
Z
A
HWEC
Z
A
BMEC
Z

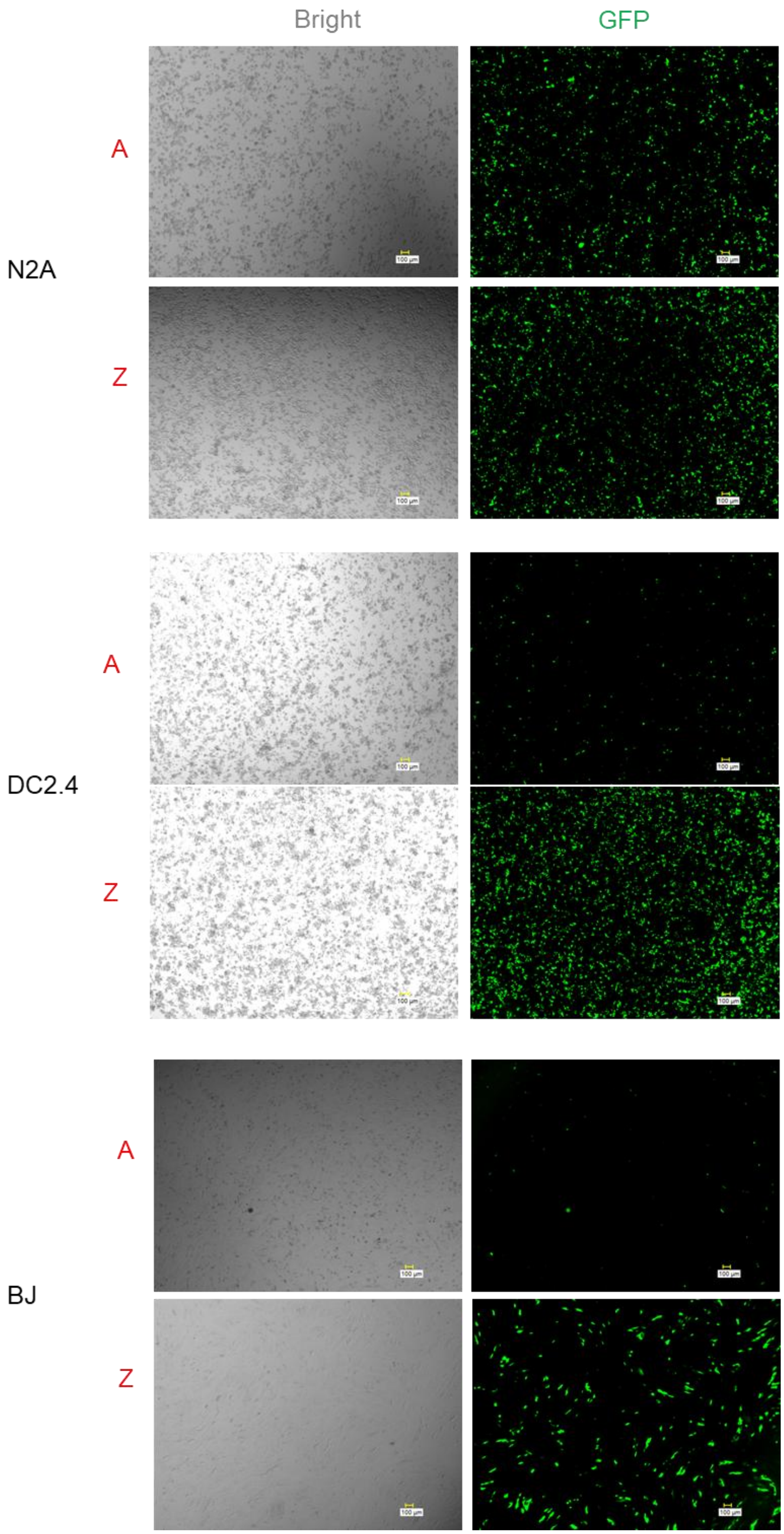

Bright
GFP
N2A
A
Z
DC2.4
A
Z
BJ
A
Z
100 μm

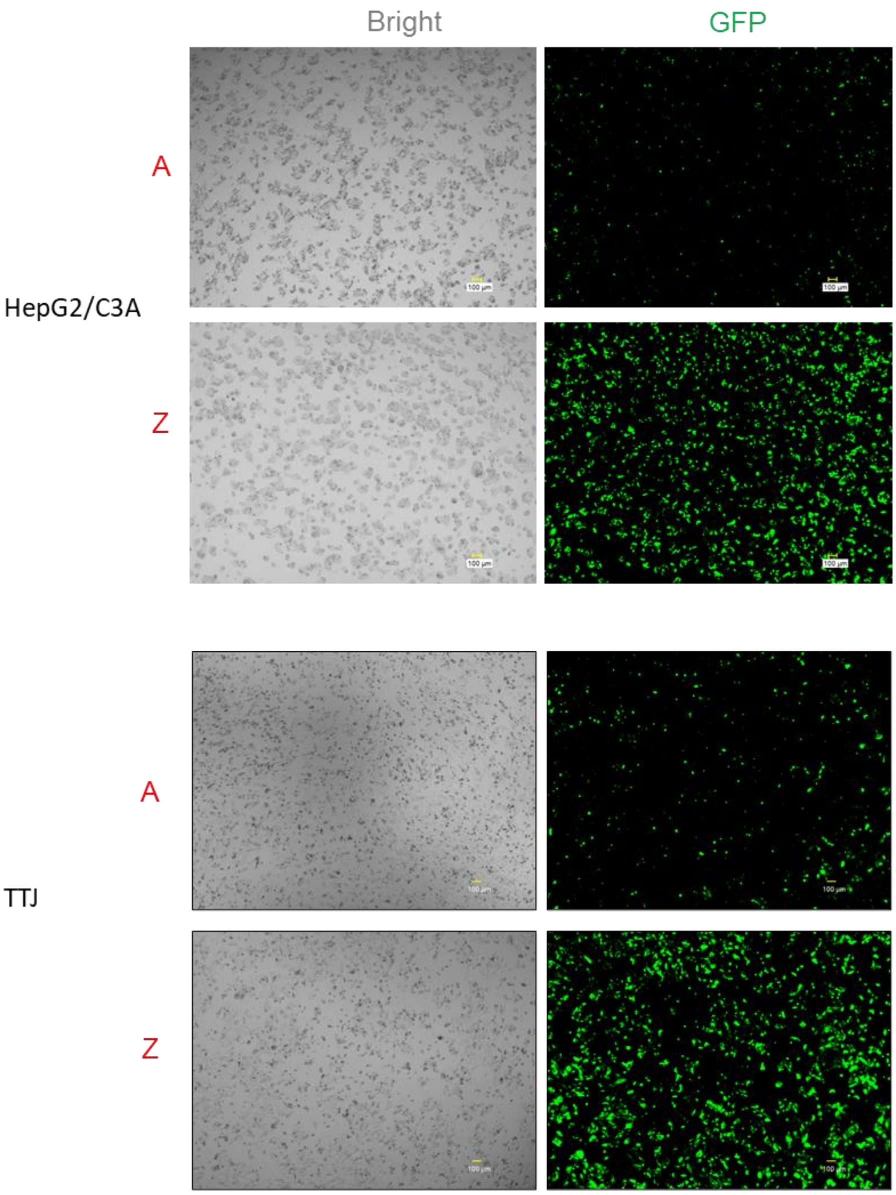


**Fig. s30 Representative brightfield and GFP images of transfected cells in Fig. s29.**

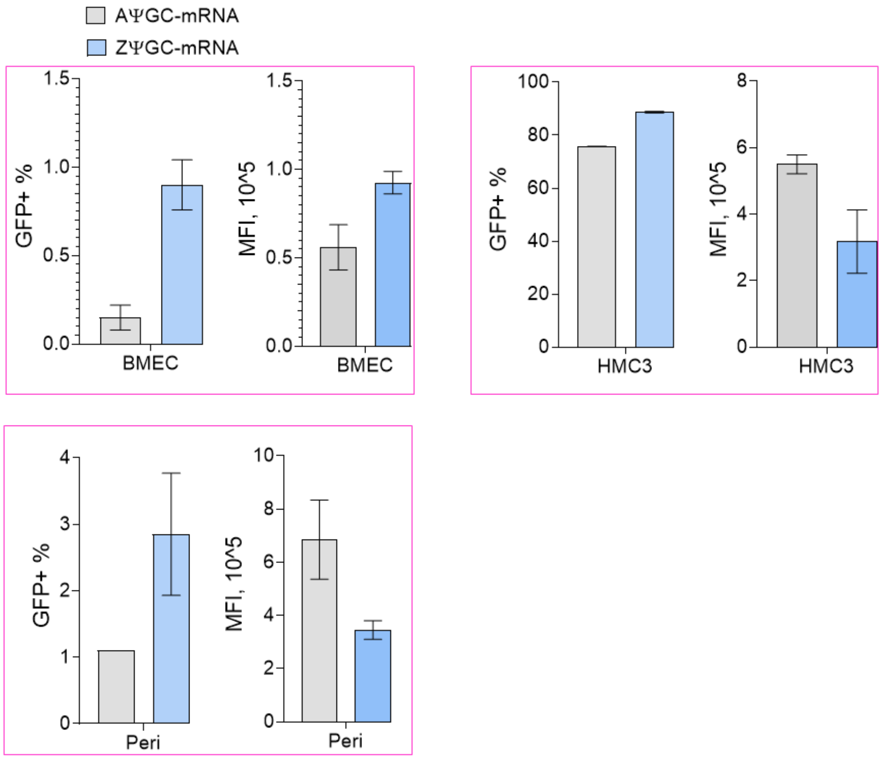


**Fig. s31 Expression of EGFP mRNA written by AΨGC and ZΨGC in neural cell lines.** Data are presented as mean ± SD (n = 3).

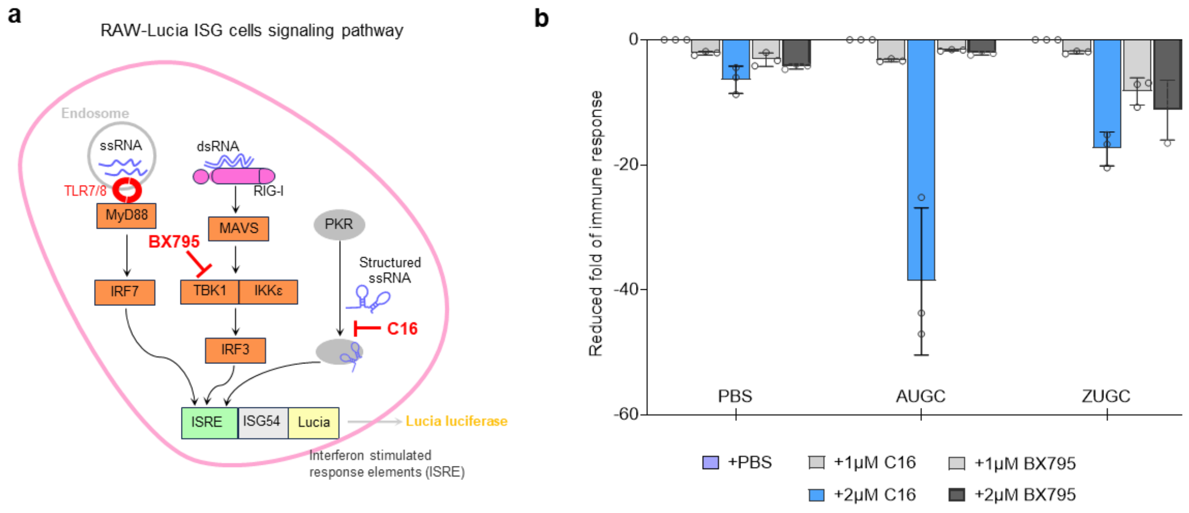


**Fig. s32 Exogenous IVT RNA sensing pathway in ISG cells.** (**a**) Schematic illustration of C16 and BX795 inhibitor targets. (**b**) Fold reduction in immune responses indued by lipofection of mRNA with or without inhibitor treatment. Data are presented as mean ± SD (n = 3). Statistical analysis was performed using 2-way ANOVA with Tukey's multiple comparisons test. *P* values of <0.05 were considered statistically significant. ns, not significant.

**Table 1 Templates sequences for long RNA synthesis**

| |
|---|
| EGFP-UTR1 sequence A, 24.1%; C, 32.3%; G, 27.9%; T, 15.7%.<br><br>GCCTAGGCTTTTGCAAAAAGCTCCCGGGAGCTTGTATATCCATTTTCGGATCTGATCAGCACGTGTGC<br>CACCATGGTGAGCAAGGGCGAGGAGCTGTTCACCGGGGTGGTGCCCATCCTGGTCGAGCTGGACGG<br>CGACGTAAACGGCCACAAGTTCAGCGTGTCCGGCGAGGGCGAGGGCGATGCCACCTACGGCAAGCT<br>GACCCTGAAGTTCATCTGCACCACCGGCAAGCTGCCCGTGCCCTGGCCCACCCTCGTGACCACCCTG<br>ACCTACGGCGTGCAGTGCTTCAGCCGCTACCCCGACCACATGAAGCAGCACGACTTCTTCAAGTCCG<br>CCATGCCCGAAGGCTACGTCCAGGAGCGCACCATCTTCTTCAAGGACGACGGCAACTACAAGACCC<br>GCGCCGAGGTGAAGTTCGAGGGCGACACCCTGGTGAACCGCATCGAGCTGAAGGGCATCGACTTCA<br>AGGAGGACGGCAACATCCTGGGGCACAAGCTGGAGTACAACTACAACAGCCACAACGTCTATATCAT<br>GGCCGACAAGCAGAAGAACGGCATCAAGGTGAACTTCAAGATCCGCCACAACATCGAGGACGGCAG<br>CGTGCAGCTCGCCGACCACTACCAGCAGAACACCCCCATCGGCGACGGCCCCGTGCTGCTGCCCGA<br>CAACCACTACCTGAGCACCCAGTCCGCCCTGAGCAAAGACCCCAACGAGAAGCGCGATCACATGGT<br>CCTGCTGGAGTTCGTGACCGCCGCCGGGATCACTCTCGGCATGGACGAGCTGTACAAGTAGGGAATT<br>AGATCACTAGTCCAGTCGGGGGCTAGCAAAATCAGCCTCGACTGTGCCTTCTAGTTGCCAGCCATCTG<br>TTGTTTGCCCCTCCCCCGTGCCTTCCTTGACCCTGGAAGGTGCCACTCCCACTGTCCTTTCCTAATAA<br>AATGAGGAAATTGCATCACAACACTCAACC |
| EGFP-UTR2 sequence A, 22.6%; C, 32.8%; G, 26.5%; T, 18.1%.<br><br>GCACTCGCGCTGCCATCACTCTTCCGCCGTCTTCGCCGCCATCCTCGGCGCGACTCGCTTCTTTCGGT<br>TCTACCAGGTAGAGTCCGCCGCCATCCTCCACCCAACAACTTGTCTCGCTCCGGGGAACGCTCGGAA<br>ACTCCCGGCCGCCGCCACCCGCGTCTGTTCTGTTACACAAGGGAAGAAAAGCCGCTGCCGCACTCC<br>GAGTGTGCCACCATGGTGAGCAAGGGCGAGGAGCTGTTCACCGGGGTGGTGCCCATCCTGGTCGAG<br>CTGGACGGCGACGTAAACGGCCACAAGTTCAGCGTGTCCGGCGAGGGCGAGGGCGATGCCACCTAC<br>GGCAAGCTGACCCTGAAGTTCATCTGCACCACCGGCAAGCTGCCCGTGCCCTGGCCCACCCTCGTGA<br>CCACCCTGACCTACGGCGTGCAGTGCTTCAGCCGCTACCCCGACCACATGAAGCAGCACGACTTCTT<br>CAAGTCCGCCATGCCCGAAGGCTACGTCCAGGAGCGCACCATCTTCTTCAAGGACGACGGCAACTAC<br>AAGACCCGCGCCGAGGTGAAGTTCGAGGGCGACACCCTGGTGAACCGCATCGAGCTGAAGGGCATC<br>GACTTCAAGGAGGACGGCAACATCCTGGGGCACAAGCTGGAGTACAACTACAACAGCCACAACGTC<br>TATATCATGGCCGACAAGCAGAAGAACGGCATCAAGGTGAACTTCAAGATCCGCCACAACATCGAGG<br>ACGGCAGCGTGCAGCTCGCCGACCACTACCAGCAGAACACCCCCATCGGCGACGGCCCCGTGCTGC<br>TGCCCGACAACCACTACCTGAGCACCCAGTCCGCCCTGAGCAAAGACCCCAACGAGAAGCGCGATC<br>ACATGGTCCTGCTGGAGTTCGTGACCGCCGCCGGGATCACTCTCGGCATGGACGAGCTGTACAAGTA<br>AAGCGGCCGCACTCCTCAGGTGCAGGCTGCCTATCAGAAGGTGGTGGCTGGTGTGGCCAATGCCCTG<br>GCTCACAAATACCACTGAGATCTTTTTCCCTCTGCCAAAAATTATGGGGACATCATGAAGCCCCTTGA<br>GCATCTGACTTCTGGCTAATAAAGGAAATTTATTTTCATTGCAATAGTGTGTTGGAATTTTTTGTGTCTC<br>TCA |
| Cre-UTR2 sequence A, 24%; C, 25.6%; G, 26.5%; T, 24%.<br><br>GCACTCGCGCTGCCATCACTCTTCCGCCGTCTTCGCCGCCATCCTCGGCGCGACTCGCTTCTTTCGGT<br>TCTACCAGGTAGAGTCCGCCGCCATCCTCCACCCAACAACTTGTCTCGCTCCGGGGAACGCTCGGAA<br>ACTCCCGGCCGCCGCCACCCGCGTCTGTTCTGTTACACAAGGGAAGAAAAGCCGCTGCCGCACTCC<br>GAGTGTCGCCACCATGCCCAAGAAGAAGAGGAAGGTGTCCAATTTACTGACCGTACACCAAAATTTG<br>CCTGCATTACCCGTCGATGCAACGAGTGATGAGGTTCGCAAGAACCTGATGGACATGTTCAGGGATC<br>GCCAGGCGTTTTCTGAGCATACCTGGAAAATGCTTCAGTCCGTTTGCCGGTCGTGGGCGGCATGGTG<br>CAAGTTGAATAACCGGAAATGGTTTCCCGCAGAACCTGAAGATGTTCGCGATTATCTTCTATATCTTCA<br>GGCGCGCGGTCTGGCAGTAAAAACTATCCAGCAACATTTGGGCCAGCTAAACATGCTTCATCGTCGG<br>TCCGGGCTGCCACGACCAAGTGACAGCAATGCTGTTTCACTGGTTATGCGGCGGATCCGAAAAGAAA<br>ACGTTGATGCCGGTGAACGTGCAAAACAGGCTCTAGCGTTCGAACGCACTGATTTCGACCAGGTTCG<br>TTCACTCATGGAAAATAGCGATCGCTGCCAGGATATACGTAATCTGGCATTTCTGGGGATTGCTTATAA<br>CACCCTGTTACGTATAGCCGAAATTGCCAGGATCAGGGTTAAAGATATCTCACGTACTGACGGTGGGA<br>GAATGTTAATCCATATTGGCAGAACGAAAACGCTGGTTAGCACCGCAGGTGTAGAGAAGGCACTTAG<br>CCTGGGGGTAACTAAACTGGTCGAGCGATGGATTTCCGTCTCTGGTGTAGCTGATGATCCGAATAACT |

ACCTGTTTTGCCGGGTCAGAAAAAATGGTGTTGCCGCGCCATCTGCCACCAGCCAGCTATCAACTCG
CGCCCTGGAAGGGATTTTTGAAGCAACTCATCGATTGATTTACGGCGCTAAGGATGACTCTGGTCAGA
GATACCTGGCCTGGTCTGGACACAGTGCCCGTGTCGGAGCCGCGCGAGATATGGCCCGCGCTGGAGT
TTCAATACCGGAGATCATGCAAGCTGGTGGCTGGACCAATGTAAATATTGTCATGAACTATATCCGTAA
CCTGGATAGTGAAACAGGGGCAATGGTGCGCCTGCTGGAAGATGGCGATTAGAGCGGCCGCACTCCT
CAGGTGCAGGCTGCCTATCAGAAGGTGGTGGCTGGTGTGGCCAATGCCCTGGCTCACAAATACCACT
GAGATCTTTTTCCCTCTGCCAAAAATTATGGGGACATCATGAAGCCCCTTGAGCATCTGACTTCTGGC
TAATAAAGGAAATTTATTTTCATTGCAATAGTGTGTTGGAATTTTTTGTGTCTCTCACTCGGAAGGACA

Cre-UTR1 sequence A, 25.1%; C, 23.2%; G, 26.9%; T, 24.8%.

GGCCTAGGCTTTTGCAAAAAGCTCCCGGGAGCTTGTATATCCATTTTCGGATCTGATCAGCACGTGTG
CCACCATGCCCAAGAAGAAGAGGAAGGTGTCCAATTTACTGACCGTACACCAAAATTTGCCTGCATT
ACCCGTCGATGCAACGAGTGATGAGGTTCGCAAGAACCTGATGGACATGTTCAGGGATCGCCAGGCG
TTTTCTGAGCATACCTGGAAAATGCTTCAGTCCGTTTGCCGGTCGTGGGCGGCATGGTGCAAGTTGA
ATAACCGGAAATGGTTTCCCGCAGAACCTGAAGATGTTCGCGATTATCTTCTATATCTTCAGGCGCGC
GGTCTGGCAGTAAAAACTATCCAGCAACATTTGGGCCAGCTAAACATGCTTCATCGTCGGTCCGGGCT
GCCACGACCAAGTGACAGCAATGCTGTTTCACTGGTTATGCGGCGGATCCGAAAAGAAAACGTTGAT
GCCGGTGAACGTGCAAAACAGGCTCTAGCGTTCGAACGCACTGATTTCGACCAGGTTCGTTCACTCA
TGGAAAATAGCGATCGCTGCCAGGATATACGTAATCTGGCATTTCTGGGGATTGCTTATAACACCCTGT
TACGTATAGCCGAAATTGCCAGGATCAGGGTTAAAGATATCTCACGTACTGACGGTGGGAGAATGTTA
ATCCATATTGGCAGAACGAAAACGCTGGTTAGCACCGCAGGTGTAGAGAAGGCACTTAGCCTGGGGG
TAACTAAACTGGTCGAGCGATGGATTTCCGTCTCTGGTGTAGCTGATGATCCGAATAACTACCTGTTTT
GCCGGGTCAGAAAAAATGGTGTTGCCGCGCCATCTGCCACCAGCCAGCTATCAACTCGCGCCCTGGA
AGGGATTTTTGAAGCAACTCATCGATTGATTTACGGCGCTAAGGATGACTCTGGTCAGAGATACCTGG
CCTGGTCTGGACACAGTGCCCGTGTCGGAGCCGCGCGAGATATGGCCCGCGCTGGAGTTTCAATACC
GGAGATCATGCAAGCTGGTGGCTGGACCAATGTAAATATTGTCATGAACTATATCCGTAACCTGGATA
GTGAAACAGGGGCAATGGTGCGCCTGCTGGAAGATGGCGATTAGAGCGGCCGCACTCCTCAGGTGC
AGGCTGCCTATCAGAAGGTGGTGGCTGGTGTGGCCAATGCCCTGGCTCACAAATACCACTGAGATCT
TTTTCCCTCTGCCAAAAATTATGGGGACATCATGAAGCCCCTTGAGCATCTGACTTCTGGCTAATAAA
GGAAATTTATTTTCATTGCAATAGTGTGTTGGAATTTTTTGTGTCTCTCACTCGGAAGGACA

UTR1+Fluc (reduce A)extraction A, 22.2%; C, 29.5%; G, 28.2%; T, 20.2%.

GCCTAGGCTTTTGCAAAAAGCTCCCGGGAGCTTGTATATCCATTTTCGGATCTGATCAGCACGTGTGC
CACCATGCACCACCACCACCACCACGAGGATGCCAAGAACATCAAGAAGGGCCCCGCCCCTTTCTAC
CCCCTGGAGGATGGCACAGCTGGAGAGCAGCTGCATAAGGCCATGAAGAGATACGCCCTGGTCCCTG
GCACCATCGCCTTCACCGACGCCCACATCGAGGTGGATATCACCTACGCCGAGTACTTCGAGATGTCC
GTGCGGCTGGCCGAGGCCATGAAGCGGTACGGCCTCAACACCAACCACCGGATCGTGGTTTGTAGC
GAGAACAGCCTGCAGTTCTTCATGCCTGTGCTGGGAGCCCTGTTCATCGGCGTGGCTGTGGCCCCTG
CTAATGACATCTACAACGAGCGGGAGCTGCTGAACAGCATGGGCATCAGCCAGCCTACAGTGGTGTT
CGTGTCCAAGAAGGGCCTGCAGAAGATTCTGAACGTGCAGAAGAAGCTGCCTATCATTCAGAAGATT
ATCATCATGGACTCTAAGACCGACTACCAGGGCTTTCAGAGCATGTACACCTTCGTGACAAGCCACCT
GCCACCTGGCTTCAACGAGTACGACTTCGTGCCTGAGTCTTTCGATAGAGACAAAACTATCGCCCTG
ATCATGAACAGCAGCGGATCTACAGGCCTGCCCAAGGGCGTGGCTCTGCCCCACCGGACCGCCTGCG
TGCGGTTTAGCCACGCCAGAGATCCTATCTTCGGCAATCAGATCATCCCCGACACCGCCATCCTGTCC
GTGGTGCCTTTCCACCACGGCTTCGGCATGTTCACCACCCTGGGATATCTGATCTGCGGCTTCCGCGT
GGTGCTGATGTATCGTTTCGAGGAGGAGTTGTTTCTGCGGAGCCTGCAGGACTACAAGATCCAGTCC
GCCCTGCTGGTGCCCACCCTGTTTAGCTTCTTCGCCAAGAGCACCCTGATCGACAAGTACGACCTGT
CCAACCTGCACGAGATCGCTTCTGGCGGCGCCCCTCTGTCTAAGGAGGTCGGCGAGGCCGTGGCCA
AGAGGTTCCACCTGCCTGGCATCAGACAGGGCTACGGCCTTACAGAGACAACCAGCGCCATCCTGAT
TACCCCTGAAGGCGACGACAAGCCCGGCGCCGTGGGTAAGGTGGTCCCTTTTTTTGAGGCCAAGGT
GGTGGACCTGGATACCGGCAAGACCCTGGGCGTGAATCAAAGAGGCGAACTGTGCGTGCGCGGCCC
TATGATCATGAGCGGCTATGTGAACAACCCTGAGGCCACCAATGCCCTGATCGACAAGGACGGCTGG
CTGCACAGCGGAGATATCGCCTACTGGGACGAGGATGAACATTTCTTCATCGTGGATAGACTGAAGTC
TCTGATCAAGTACAAGGGATACCAAGTGGCCCCTGCTGAACTGGAGAGCATCCTGCTGCAGCACCCT
AACATCTTTGACGCCGGAGTGGCCGGACTTCCAGATGATGACGCCGGCGAGCTGCCCGCTGCTGTTG

| |
|---|
| TGGTCCTGGAACACGGCAAGACCATGACCGAGAAGGAGATCGTGGACTACGTGGCCAGCCAGGTGA<br>CCACAGCCAAGAAGCTGCGGGGCGGTGTGGTGTTCGTGGACGAAGTGCCAAAGGGCCTGACCGGC<br>AAGCTGGACGCCAGGAAGATCCGGGAGATCCTGATCAAGGCCAAGAAGGGCGGCAAGATCGCCGTG<br>GATTACAAAGACGATGACGATAAGTGATTAATTAAGCTGCCTTCTGCGGGGCTTGCCTTCTGGCCATG<br>CCCTTCTTCTCTCCCTTGCACCTGTACCTCTTGGTCTTTGAATAAAGCCTGAGTAGG |
| UTR2-Fluc A, 27.4%; C, 22.1%; G, 24.7%; T, 25.8%.<br><br>GCACTCGCGCTGCCATCACTCTTCCGCCGTCTTCGCCGCCATCCTCGGCGCGACTCGCTTCTTTCGGT<br>TCTACCAGGTAGAGTCCGCCGCCATCCTCCACCCAACAACTTGTCTCGCTCCGGGGAACGCTCGGAA<br>ACTCCCGGCCGCCGCCACCCGCGTCTGTTCTGTTACACAAGGGAAGAAAAGCCGCTGCCGCACTCC<br>GAGTGTCGCCACCATGGAAGACGCCAAAAACATAAAGAAAGGCCCGGCGCCATTCTATCCGCTGGA<br>AGATGGAACCGCTGGAGAGCAACTGCATAAGGCTATGAAGAGATACGCCCTGGTTCCTGGAACAATT<br>GCTTTTACAGATGCACATATCGAGGTGGACATCACTTACGCTGAGTACTTCGAAATGTCCGTTCGGTT<br>GGCAGAAGCTATGAAACGATATGGGCTGAATACAAATCACAGAATCGTCGTATGCAGTGAAAACTCT<br>CTTCAATTCTTTATGCCGGTGTTGGGCGCGTTATTTATCGGAGTTGCAGTTGCGCCCGCGAACGACATT<br>TATAATGAACGTGAATTGCTCAACAGTATGGGCATTTCGCAGCCTACCGTGGTGTTCGTTTCCAAAAA<br>GGGGTTGCAAAAAATTTTGAACGTGCAAAAAAAGCTCCCAATCATCCAAAAAATTATTATCATGGATT<br>CTAAAACGGATTACCAGGGATTTCAGTCGATGTACACGTTCGTCACATCTCATCTACCTCCCGGTTTTA<br>ATGAATACGATTTTGTGCCAGAGTCCTTCGATAGGGACAAGACAATTGCACTGATCATGAACTCCTCT<br>GGATCTACTGGTCTGCCTAAAGGTGTCGCTCTGCCTCATAGAACTGCCTGCGTGAGATTCTCGCATGC<br>CAGAGATCCTATTTTTGGCAATCAAATCATTCCGGATACTGCGATTTTAAGTGTTGTTCCATTCCATCAC<br>GGTTTTGGAATGTTTACTACACTCGGATATTTGATATGTGGATTTCGAGTCGTCTTAATGTATAGATTTG<br>AAGAAGAGCTGTTTCTGAGGAGCCTTCAGGATTACAAGATTCAAAGTGCGCTGCTGGTGCCAACCCT<br>ATTCTCCTTCTTCGCCAAAAGCACTCTGATTGACAAATACGATTTATCTAATTTACACGAAATTGCTTC<br>TGGTGGCGCTCCCCTCTCTAAGGAAGTCGGGGAAGCGGTTGCCAAGAGGTTCCATCTGCCAGGTATC<br>AGGCAAGGATATGGGCTCACTGAGACTACATCAGCTATTCTGATTACACCCGAGGGGGATGATAAACC<br>GGGCGCGGTCGGTAAAGTTGTTCCATTTTTTGAAGCGAAGGTTGTGGATCTGGATACCGGGAAAACG<br>CTGGGCGTTAATCAAAGAGGCGAACTGTGTGTGAGAGGTCCTATGATTATGTCCGGTTATGTAAACAA<br>TCCGGAAGCGACCAACGCCTTGATTGACAAGGATGGATGGCTACATTCTGGAGACATAGCTTACTGG<br>GACGAAGACGAACACTTCTTCATCGTTGACCGCCTGAAGTCTCTGATTAAGTACAAAGGCTATCAGG<br>TGGCTCCCGCTGAATTGGAATCCATCTTGCTCCAACACCCCAACATCTTCGACGCAGGTGTCGCAGGT<br>CTTCCCGACGATGACGCCGGTGAACTTCCCGCCGCCGTTGTTGTTTTGGAGCACGGAAAGACGATGA<br>CGGAAAAAGAGATCGTGGATTACGTCGCCAGTCAAGTAACAACCGCGAAAAAGTTGCGCGGAGGAG<br>TTGTGTTTGTGGACGAAGTACCGAAAGGTCTTACCGGAAAACTCGACGCAAGAAAAATCAGAGAGA<br>TCCTCATAAAGGCCAAGAAGGGCGGAAAGATCGCCGTGTAAAGCGGCCGCACTCCTCAGGTGCAGG<br>CTGCCTATCAGAAGGTGGTGGCTGGTGTGGCCAATGCCCTGGCTCACAAATACCACTGAGATCTTTTT<br>CCCTCTGCCAAAAATTATGGGGACATCATGAAGCCCCTTGAGCATCTGACTTCTGGCTAATAAAGGAA<br>ATTTATTTTCATTGCAATAGTGTGTTGGAATTTTTTGTGTCTCTCACTCGGAAGGACATATGGG |
| DNA template for dsRNA-sense (T7)<br>TAATACGACTCACTATAGGCCTAGGCTTTTGCAAAAAGCTCCCGGGAGCTTGTATATCCATTTTCGGAT<br>CTGATCAGCACGTGTGCCG<br><br>DNA template for dsRNA-anti sense (T7)<br>TAATACGACTCACTATA<br>GGCACACGTGCTGATCAGATCCGAAAATGGATATACAAGCTCCCGGGAGCTTTTTGCAAAAGCCTAG<br>GCC |

#### Table 2 DNA templates for Cas9 crRNA and tracrRNA

| **Name** | **Sequence (T7) TAATACGACTCACTATA+ ss** |
|---|---|
| ss1 | GTAAACAAAGCATAGACTGAGTTTTAGAGCTATGCTGTTTTG |
| ss2 | GAACATAAAGAATAGAATGAGTTTTAGAGCTATGCTGTTTTG |
| ss3 (tracrRNA) | GGGGAACCATTCAAAACAGCATAGCAAGTTAAAATAAGGCTAGTCCGTTATCAACTTGAAAAAGTGGCACCGAGTCGGTGCTTTTTTT |

#### Table 3 Chemical synthesized ssRNA

| **Name** | **Sequence** |
|---|---|
| sgRNA_SYN Genscirpt | 5'mG*mU*mA*rArArCrArArArGrCrArUrArGrArCrUrGrArGrUrUrUrUrArGrArGrCrUrArGrArArArUrArGrCrArArGrUrUrArArArArUrArArGrGrCrUrArGrUrCrCrGrUrUrArUrCrArArCrUrUrGrArArArArArGrUrGrGrCrArCrCrGrArGrUrCrGrGrUrGrCrU*mU*mU*mU-3' |
| IDT Alt-R CRISPR-Cas9 tracrRNA | 5'AGCAUAGCAAGUUAAAAUAAGGCUAGUCCGUUAUCAACUUGAAAAAGUGGCACCGAGUCGGUGCUUU-3’ Note: The modifications on this tracrRNA are proprietary and cannot be shared. |

### Table 4 DNA templates for Cas12a crRNA IVT

| Name | Sequence 2 (T7) TAATACGACTCACTATA+ cr | A count | T count | A % | T % | T/A | C count | G count | U/C ratio | U/G ratio | "RT" count |
|---|---|---|---|---|---|---|---|---|---|---|---|
| cr1 | GGAATTTCTACTGTTGTAGATAATCTGAAAATATATTAAT | 15 | 16 | 37 | 39 | 1.1 | 3 | 6 | 5.3 | 2.7 | |
| cr2 | GGAATTTCTACTGTTGTAGATTCAGATATTCTACTTTGTT | 9 | 19 | 22 | 46 | 2.1 | 5 | 7 | 3.8 | 2.7 | |
| cr3 | GGAATTTCTACTGTTGTAGATGATCTCTAAATTACATAAAC | 14 | 15 | 34 | 37 | 1.1 | 6 | 6 | 2.5 | 2.5 | |
| cr4 | GGAATTTCTACTGTTGTAGATTTTACCTTCTTCTTGATATG | 8 | 20 | 20 | 49 | 2.5 | 6 | 7 | 3.3 | 2.9 | 6.0 |
| cr5 | GGAATTTCTACTGTTGTAGATATTTTGAGGCTCTGCAAAGT | 10 | 16 | 24 | 39 | 1.6 | 5 | 10 | 3.2 | 1.6 | 6.0 |
| cr6 | GGAATTTCTACTGTTGTAGATAACTATATTTTTTCACATAG | 12 | 18 | 29 | 44 | 1.5 | 5 | 6 | 3.6 | 3.0 | 7.0 |
| cr7 | GGAATTTCTACTGTTGTAGATCCCAGAGTCCTTCAAAGATA | 12 | 13 | 29 | 32 | 1.1 | 8 | 8 | 1.6 | 1.6 | 6.0 |
| cr8 | GGAATTTCTACTGTTGTAGATACATAATAATTTAAAATATC | 16 | 16 | 39 | 39 | 1.0 | 4 | 5 | 4.0 | 3.2 | 9.0 |
| cr9 | GGAATTTCTACTGTTGTAGATAAATATCCTTATTCCATTAT | 12 | 18 | 29 | 44 | 1.5 | 6 | 5 | 3.0 | 3.6 | 9.0 |
| cr10 | GGAATTTCTACTGTTGTAGATAGTTGTAGATATGATTTAGC | 11 | 17 | 27 | 41 | 1.5 | 3 | 10 | 5.7 | 1.7 | 9.0 |
| cr11 | GGAATTTCTACTGTTGTAGATGCTCACAGCATACATATATA | 13 | 14 | 32 | 34 | 1.1 | 7 | 7 | 2.0 | 2.0 | 8.0 |
| cr12 | GGAATTTCTACTGTTGTAGATATTTCTGCCATATTTCCTTC | 8 | 19 | 20 | 46 | 2.4 | 8 | 6 | 2.4 | 3.2 | 7.0 |
| cr13 | GGAATTTCTACTGTTGTAGATAATTATTATGTTAAAATGTT | 13 | 19 | 32 | 46 | 1.5 | 2 | 7 | 9.5 | 2.7 | 10.0 |
| cr14 | GGAATTTCTACTGTTGTAGATTACAGACAATCCAAGAAGGT | 14 | 12 | 34 | 29 | 0.9 | 6 | 9 | 2.0 | 1.3 | 6.0 |
| cr15 | GGAATTTCTACTGTTGTAGATAAGGCCAAACCTCGGCTTAC | 11 | 12 | 27 | 29 | 1.1 | 9 | 9 | 1.3 | 1.3 | 4.0 |
| cr16 | GGAATTTCTACTGTTGTAGATTTCATATGTTCTTCTAGCTT | 8 | 20 | 20 | 49 | 2.5 | 6 | 7 | 3.3 | 2.9 | 7.0 |
| cr17 | GGAATTTCTACTGTTGTAGATCACTAACACGCATATTTGAT | 12 | 15 | 29 | 37 | 1.3 | 7 | 7 | 2.1 | 2.1 | 7.0 |
| cr18 | GGAATTTCTACTGTTGTAGATTGTAATTTAGAGATCTAAAC | 13 | 16 | 32 | 39 | 1.2 | 4 | 8 | 4.0 | 2.0 | 7.0 |
| cr19 | GGAATTTCTACTGTTGTAGATGAGATCTAAACAAAGTAGAA | 16 | 12 | 39 | 29 | 0.8 | 4 | 9 | 3.0 | 1.3 | 6.0 |
| cr20 | GGAATTTCTACTGTTGTAGATGACAGAAATTTAGAAGTCAT | 14 | 14 | 34 | 34 | 1.0 | 4 | 9 | 3.5 | 1.6 | 7.0 |
| cr21 | GGAATTTCTACTGTTGTAGATGAAGTCATAATTTTACTGAA | 13 | 16 | 32 | 39 | 1.2 | 4 | 8 | 4.0 | 2.0 | 7.0 |
| cr22 | GGAATTTCTACTGTTGTAGATCTGAATATCTATGCATTAAT | 12 | 17 | 29 | 41 | 1.4 | 5 | 7 | 3.4 | 2.4 | 9.0 |
| cr23 | GGAATTTCTACTGTTGTAGATTTATATATGTGATACATTTT | 11 | 20 | 27 | 49 | 1.8 | 3 | 7 | 6.7 | 2.9 | 10.0 |
| cr24 | GGAATTTCTACTGTTGTAGATACTTACCTACAGTGAAACAT | 13 | 14 | 32 | 34 | 1.1 | 7 | 7 | 2.0 | 2.0 | 6.0 |
| cr25 | GGAATTTCTACTGTTGTAGATGGCAATATATGATTAATATA | 14 | 16 | 34 | 39 | 1.1 | 3 | 8 | 5.3 | 2.0 | 10.0 |
| cr26 | GGAATTTCTACTGTTGTAGATAAAAGAATAAGTATACAACA | 18 | 12 | 44 | 29 | 0.7 | 4 | 7 | 3.0 | 1.7 | 7.0 |
| cr27 | GGAATTTCTACTGTTGTAGATTATGTCCTCACATCACAGAA | 12 | 14 | 29 | 34 | 1.2 | 8 | 7 | 1.8 | 2.0 | 7.0 |

**Table 5 Primers for PCR**

| | |
|---|---|
| HG-IVT template PCR-F | ttgttgATTTAGGTGACACTATAGcctaggcttttgcaaaaagctcc |
| HG-IVT template PCR-R | ggttgagtgttgtgatgcaatttcc |
| | *Amplify EGFP-UTR1 template using this pair.* |
| G300, SP6-CoNeoUTR2-3-F: | TTGTTGATTTAGGTGACACTATA GCACTCGCGCTGCCATCACTCTT |
| | *Paired with G301 primer to amplify template p94, p157, p158 plasmids.* |
| G320 | TTGTTGATTTAGGTGACACTATA GG CACTCGCGCTGCCATCACTCTT |
| | *Generated from modified G300, SP6 promoter, GG start site* |
| G301 beta-globin-R: | TGTCCTTCCGAGTGAGAGACA |
| | For template p94,p157,p158 |
| G324 | CCTAGGCTTTTGCAAAAAGCTCCCGGGAGCTTGTATATCCATTTTCGGATCTGATCAGC ACGTGTGCCACC atggaagacgccaaaaacataaag |
| | *Paired with G301 primer, amplify FLuc using P157 plasmid.* |
| G325 | CCTAGGCTTTTGCAAAAAGCTCCCGGGAGCTTGTATATCCATTTTCGGATCTGATCAGC ACGTGTGCCACC ATGCCCAAGAAGAAGAGGAAGGT |
| | *Paired with G301 primer, amplify Cre using P158 plasmid.* |
| G323 | ttgttgATTTAGGTGACACTATA GG cctaggcttttgcaaaaagctcc |
| | *Paired with G301 primer, amplify PCR fragment (G324/G301) using fragments carrying UTR1.* |
| G312 | TTGTTGATTTAGGTGACACTATA GG aaataagagagaaaagaagagtaagaagaaatataag |
| G313 | CTTCCTACTCAGGCTTTATTCAAAGACCA |
| | *Amplify ABE from P228 or P261 plasmids using pairs G312/G313 for SP6 IVT.* |
